# Optimal cues for transmission investment in malaria parasites

**DOI:** 10.1101/2025.02.26.640285

**Authors:** Avril L. Wang, Megan A. Greischar, Nicole Mideo

## Abstract

The timing of investment into reproduction is a key determinant of lifetime reproductive success (fitness). Many organisms exhibit plastic, i.e., environmentally-responsive, investment strategies, raising the questions of what environmental cues trigger responses and why organisms have evolved to respond to those particular cues. For malaria parasites (*Plasmodium spp*.), investment into the production of specialized transmission stages (versus stages that replicate asexually within the host) is synonymous with reproductive investment and also plastic, responding to host- and parasite-derived factors. Previous theory has identified optimal plastic transmission investment strategies for the rodent malaria parasite, *Plasmodium chabaudi*, as a function of the time since infection, implicitly assuming that parasites have perfect information about the within-host environment and how it is changing. We extend that theory to ask which cue(s) *should* parasites use? Put another way, which cue(s) maximize parasite fitness, quantified as host infectiousness during acute infection? Our results show that sensing a parasite-associated cue, e.g., the abundance of infected red blood cells or transmission stages, allows parasites to achieve fitness approaching that of the optimal time-varying strategy, but only when parasites perceive the cue non-linearly, responding more sensitively to changes at low densities. However, no single cue can recreate the best time-varying strategy or allow parasites to adopt terminal investment as the infection ends, a classic expectation for reproductive investment. Sensing two cues—log-transformed infected and uninfected red blood cell abundance—enables parasites to accurately track the progression of the infection, permits terminal investment, and recovers the fitness of the optimal time-varying investment strategy. Importantly, parasites that detect two cues more efficiently exploit hosts, resulting in higher virulence compared with those sensing only one cue. Finally, our results suggest a potential tradeoff between achieving an optimal transmission investment strategy in a given host environment and robustness in the face of environmental or developmental fluctuations.

## Introduction

A key goal of life history theory is to understand how organisms should invest in reproduction over the course of their lives and explaining the substantial variation in patterns observed (e.g., [1]). Reproductive investment entails a resource allocation tradeoff: resources allocated to growth cannot, generally, also be allocated to reproduction. Classic work predicts that reproductive investment will increase with age [2, 3]. Additionally, reproductive investment is expected to depend on the state of an organism, a product of intrinsic, physiological factors and external, environmental ones [4]. Experimental data have shown that investment can be plastic, with age and state both shaping an organism’s reproductive efforts to maximize lifetime fitness (e.g., [5, 6]). How organisms estimate the time remaining for reproduction—that is, what cues they utilize from their own physiology and the environment—remains an open question for many systems.

For sexually-reproducing parasites, patterns of reproductive investment can have clinical and epidemiological consequences. Protozoan parasites of *Plasmodium* spp. —the etiological agents of malaria —infect red blood cells of their host, and a given infected cell will either produce more asexual, cell-infecting stages (“growth” of an infection) or produce a single gametocyte, a sexually-differentiated stage that can be transmitted to mosquito vectors (potentially “reproducing” an infection). The proportion of infected cells that produce transmissible gametocytes, known in the malaria literature as the “conversion rate,” is a measure of investment in reproduction/transmission [7]. This trait has a direct link to infectiousness, since more gametocytes mean greater chance of onward transmission to a biting mosquito [8–10] and can influence the virulence or severity of infection, since all else equal, lower transmission investment means more host-damaging asexual parasites are produced [7].

Importantly, transmission investment in *Plasmodium* spp. is plastic, changing in response to conspecific density [11, 12], replication rate [13], parasite-conditioned media [14, 15] (which could contain infected red blood cell (iRBC)-derived vesicles [16]), RBC age structure/availability [13, 17–19], and antimalarial exposure (e.g., [13, 20]). Understanding why transmission investment is plastic and what the optimal plastic strategy is has been the subject of several prior studies [7, 13, 21–24]. In one example, using the experimentally tractable rodent malaria system, *Plasmodium chabaudi*, and different doses of antimalarial drugs, Schneider *et al*. [13] showed that intermediate reductions in parasite density led to reduced investment in transmission while strong reductions had the opposite effect. These results are in line with standard predictions from life history theory for malaria parasites, where mild stressors should promote reproductive restraint and strong stressors that signal imminent clearance of an infection should lead to complete investment in transmission [7], i.e., terminal investment [25]. But “stress” will vary over the course of infection—even in the absence of drug treatment— as parasites deplete host resources and hosts respond via immunity. These dynamics are thought to explain why transmission investment in malaria parasites is variable over the lifespan of infections (i.e., plastic) [11–13, 22, 26, 27].

Mathematical modelling provides another route to understanding and predicting adaptive plasticity in parasite traits. Models assessing the fitness consequences of different patterns of transmission investment demonstrate that the best strategy involves initial reproductive restraint that facilitates within-host parasite population growth (rather than early investment in gametocytes that would have very low likelihood of onward transmission due to low densities), followed by an increase in investment once parasite densities have risen, culminating in terminal investment as the infection is ending [21, 23]. This work derived the optimal transmission investment strategy by assuming that investment is a function of infection age, about which parasites have perfect knowledge. In addition to assuming that parasites can, in effect, “tell time,” this work also assumes all infections proceed identically, a limitation common to models of age-specific reproductive investment [28]. In reality, time would be a poor proxy for environmental conditions since the pace of infection changes with parasite dose [29, 30], host immunity [30, 31], and myriad other factors. Therefore, understanding the links between transmission investment and within-host environmental cues will result in more biologically relevant models that can inform empirical observation of *Plasmodium spp*. transmission investment. Currently, it is unclear whether observed deviations from the theoretical optimal strategy are due to faulty theories or limitations in the cues that parasites perceive. Understanding the fitness consequences of sensing different cues can delineate those possibilities. Additionally, understanding what cues parasites should sense will enable better predictions on the influence of novel perturbations, like vaccination, on infection dynamics and outcomes.

In this study, we extend a previously-published model of within-host rodent malaria infection dynamics [23] to include transmission investment as a function of environmental cues, i.e., components of state. In our model, different cues may give rise to different early patterns of transmission investment, which will alter the trajectory of infections and feedback to influence the subsequent expression of the trait. We use numerical optimization to investigate which within-host environmental cue or cues malaria parasites “should” sense for determining transmission investment (i.e., which cues give rise to the highest parasite fitness) and the optimal shape of the relationship between each cue and transmission investment (i.e., the optimal reaction norms). We assess the consequences of perceiving different cues on infection dynamics and disease severity and quantify the robustness of different transmission investment strategies in the face of environmental and developmental stochasticity. Collectively, our study further characterizes plasticity in *P. chabaudi* life history, identifies the cues we expect parasites to be using, and introduces a mathematical framework that explicitly incorporates cue perception that could be used to generate testable hypotheses and broadly applied to other organisms.

## Results

Our mathematical model tracks within-host dynamics of infection with the rodent malaria parasite *P. chabaudi* (Fig 1A). Asexual parasites, called merozoites (*M*), invade uninfected red blood cells (RBCs; *R*). The resulting infected RBC (iRBC; *I*) can go on to produce more merozoites that are either asexually-committed (*M*; producing more merozoites if they infect an RBC) or sexually-committed (*M*_*g*_; producing a single gametocyte, *G*, if they infect an RBC). RBCs infected with these different types of merozoites are denoted as *I* and *I*_*g*_, respectively. Importantly, the proportion of iRBCs that follow these different developmental routes (*i*.*e*., producing *M*_*g*_) defines transmission investment. While previous approaches have treated transmission investment as either fixed or time-dependent (e.g., [21, 23, 32–36]), here we explicitly model it as a function of within-host environmental cues (see Materials and Methods). We select as plausible cues various measures of the density of parasites and host resources for which there is some experimental evidence to impact transmission investment in *Plasmodium* spp. (Table 1). We quantify the fitness of parasites sensing various cues using the cumulative transmission potential, which measures the summed probability of infecting mosquito vectors across a 20-day infection period, assuming a constant biting rate and mosquito abundance [23].

**Table 1.** Putative within-host environmental cues.

| Cue | Abbreviation | Experimental Evidence |
| --- | --- | --- |
| Infected RBC (iRBC) density |  | [12, 13, 16, 24] |
| Asexually-committed | I |  |
| Asexually-committed, log<sub>10</sub> | I log |  |
| Sexually-committed | I<sub>g</sub> |  |
| Sexually-committed, log<sub>10</sub> | I<sub>g</sub> log |  |
| Total (I+I<sub>g</sub>) | Total I |  |
| Total (I+I<sub>g</sub>) , log<sub>10</sub> | Total I log |  |
| Gametocyte density |  | [12, 16, 24] |
| Gametocytes | G |  |
| Gametocytes, log<sub>10</sub> | G log |  |
| Resource density |  | [12, 13, 24] |
| Uninfected RBC | R |  |
| Uninfected RBC, log<sub>10</sub> | R log |  |

**Fig 1.**
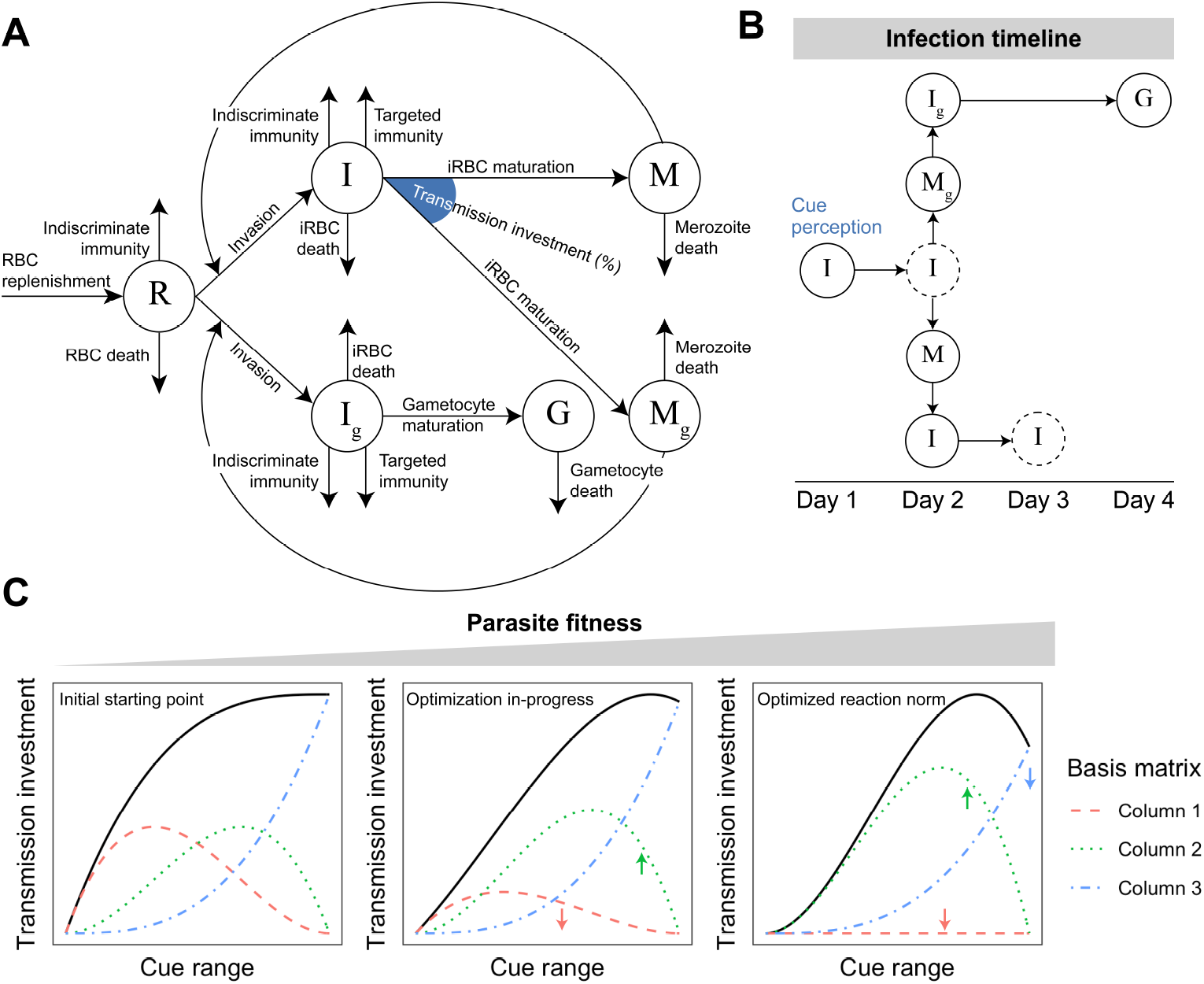
Schematic of *P. chabaudi* within-host infection dynamics and fitness optimization. (A) Flow diagram of model tracking the dynamics of red blood cells (*R*; RBCs), RBCs infected with asexual parasites (*I*), RBCs infected with sexual—but not yet mature —parasites (*I*_g_), asexually-committed merozoites (*M*), sexually-committed merozoites (*M* _g_), and gametocytes (*G*). (B) Temporal dynamics of cue perception. Cue perception occurs one day before the production of asexual/sexual merozoites, thereby introducing a delay between cue perception and the consequences of cue perception (i.e., production of more asexual infected RBCs or gametocytes). A further delay occurs in the sexual cycle since gametocytes of *P. chabaudi* take two days to mature [37]. Dashed circles indicate the bursting/maturation of a particular cohort of iRBCs. (C) Modeling reaction norm evolution. We model the relationship between the perceived cue and transmission investment (i.e., the reaction norm) as a flexible spline (black lines). Specifically, a cubic spline can specify any third order polynomial at a given x value as a linear combination of the columns in its basis matrix [38] (exemplified by the red, green, and blue dashed lines; combined they produce the solid, black line). For local optimization, an arbitrary starting spline is picked (left panel) and an optimization algorithm is used to adjust the relative weights of the basis functions until a fitness maximum is achieved (going from left to right). The small coloured arrows depict how the weight of a given basis function has changed from the previous panel.

In our model, cue perception occurs at the beginning of asexual iRBC development (Fig 1B), determining the proportion of asexual iRBCs that produces sexually-committed merozoites. Note that this modeling choice introduces a time lag between sexual commitment (as a result of cue perception) and gametocyte production, which is a departure from previous modeling work [23], but reflects empirical data on the canonical timeline of sexual commitment from the best studied system (*Plasmodium falciparum* [39]). For each putative cue, we define its associated reaction norm using flexible splines (Fig 1C). Specifically, the shape of the reaction norm is determined by a linear combination of three basis functions comprising a cubic spline. Adjusting the spline coefficients (i.e., the weights associated with each column of the spline basis matrix) changes the shape of the reaction norm. We used a combination of local and global numerical optimization algorithms (see Materials and Methods) to adjust the spline coefficients and find the reaction norm that maximizes parasite fitness over a 20-day infection, as in previous work [23, 40] (Fig 1C). We test both raw densities and logged densities as cues to expand the range of reaction norm shapes explored and because changes in infection dynamics are observed in this system when doses change by orders of magnitude [29]. For parasites sensing time, we used the same optimization process, except setting the cue range to time since infection (day 0 to 20). Finally, following [41], our within-host model includes two forms of immunity, one targeting specifically infected cells (*I* and *I*_*g*_) and one targeting all red blood cells (*I, I*_*g*_ and *R*). Thus, while resource limitation plays some role in driving infection dynamics in this model, immunity allows for the clearance of the infection.

### Cue perception mediates parasite life history and fitness

Based on previous work modeling *P. chabaudi* transmission investment as a function of time [23], we expect the following as the best strategy over a 20-day infection: parasites should delay investment early on, display a rapid increase in investment when sufficient gametocytes can be produced to ensure transmission, subtly reduce investment in the face of declining resources, and ultimately rise to the highest levels of investment shortly before the end of infection (roughly, “terminal investment” [7, 23]). Although the model we employ here has a few key differences from that work (see Materials and Methods), we find a similar optimal transmission investment pattern as in [23] when we assume that investment is a function of time (Fig 2A). When parasites instead respond to their environment and are constrained to perceive a single within-host cue, these same patterns of transmission investment cannot be fully recapitulated. Consequently, parasite fitness is lower in every case than when the cue is time (Fig 2A, left panel).

**Fig 2.**
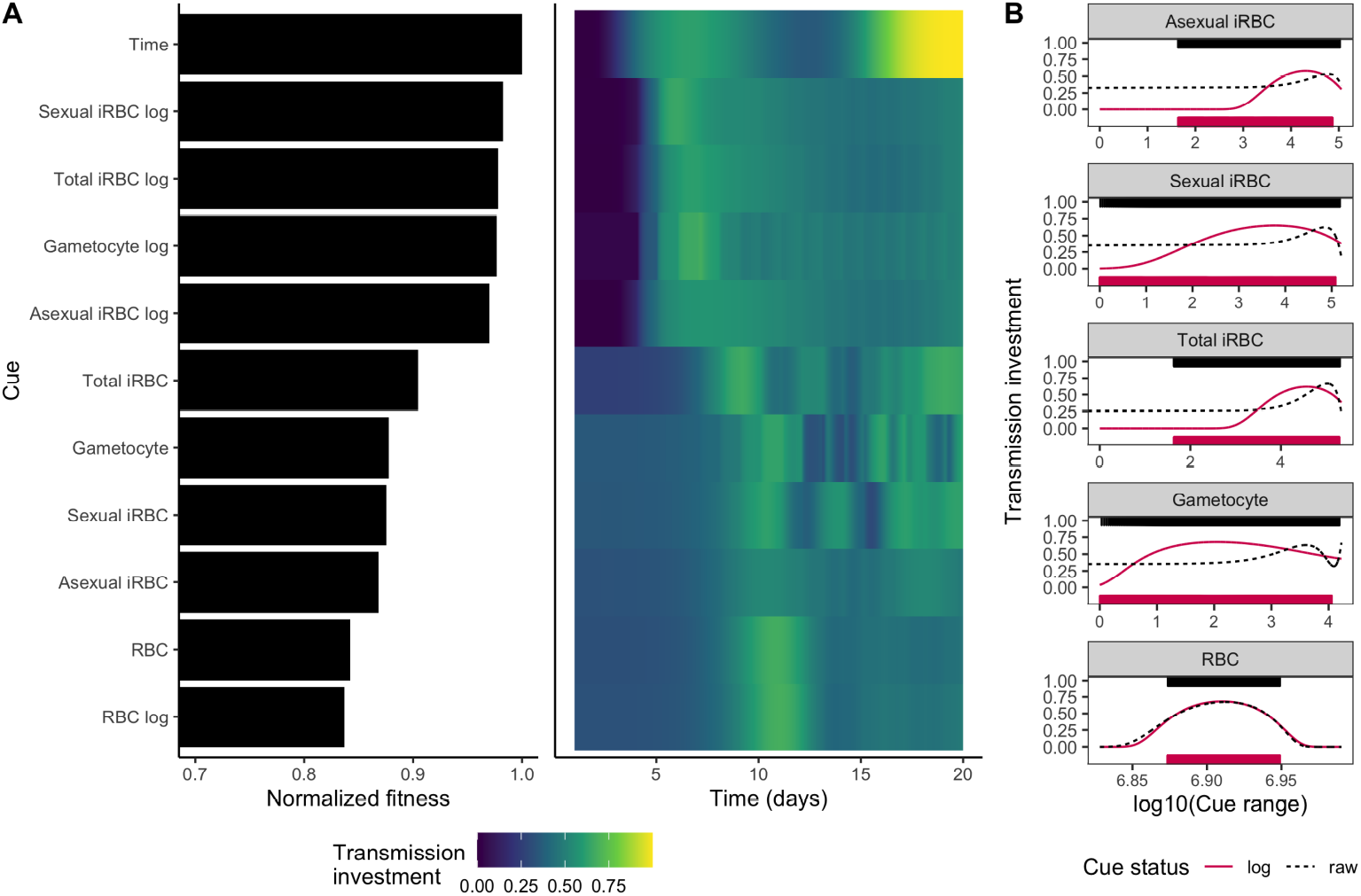
Cue perception mediates parasite fitness and dynamics of transmission investment. (A) On the left, fitness values achieved by parasites adopting the optimal transmission investment strategy for each cue, normalized with respect to the time-varying strategy, are displayed in descending order. On the right, the dynamics of transmission investment, given the optimal reaction norms, are plotted. Investment varies from dark blue (low values) to yellow (high values). The first four cues after time, which include measures of parasite density on a log_10_ scale, enable parasites to obtain the highest fitness and delay in transmission investment. (B) Optimal reaction norms for different cues. Each facet displays the reaction norms for raw (black, dotted lines) or log-transformed (red, solid lines) cues. The ranges of cue values “experienced” by a parasite adopting the optimal reaction norm are displayed above (raw) or below (log-transformed) the reaction norms as rug plots (horizontal bars). Based on the shape of the reaction norms, we conclude that sensing log-transformed parasite-centric cues enables more sensitive detection of (and responses to) the initial rise in parasite abundance.

The environmental cues that result in the highest parasite fitness include logged parasite-centric cues (e.g., log_10_ densities of sexually-committed iRBCs). Perceiving these cues is sufficient to produce the initial delay in transmission investment, but the delay is longer than parasites sensing time as the cue, and there is no increase in investment near the end of infection (Fig 2A, right panel). Given that parasite densities rise non-linearly at the start of infections (both simulated and real [11, 42–45]), sensing logged rather than raw densities results in reaction norms that are more sensitive to those early changes (Fig 2B), enabling parasites to increase transmission investment after the densities have reached a rough threshold. In contrast, sensing raw parasite densities results in reaction norms that essentially ignore changes at the low parasite densities characteristic of early infections (Fig 2B). The density of host resources (RBCs; raw or logged) is also a poor cue for initial stages of infection, since they change very little early on and, therefore, do not allow parasites to engage in delayed transmission investment (Fig 2A). Overall, our results broadly suggest that early transmission delay necessitates sensitivity to changes at low parasite density.

### Dual cue perception is necessary for terminal investment

We found that parasites sensing a single cue cannot reproduce the expected optimal transmission investment strategy, particularly the rapid increase in transmission investment during late infection (i.e., terminal investment). Achieving terminal investment presumably requires sensing a cue that provides a unique signal at the later stages of infection. Inspecting infection dynamics produced by parasites that sense time revealed that only RBC density provides such a distinctive late-stage signal (Fig 3A). Given the poor performance of this cue in early stages of infection, these results suggest that simultaneous perception of RBC and a parasite-centric cue could provide infection stage-specific information that enables both early reproductive restraint and terminal investment. Consistent with this expectation, in a rodent malaria experiment, including variables that reflect parasite density and host resources (specifically, replication rate and changes in RBC density) in statistical models explained substantially more variation in transmission investment than the parasite-centric variable alone [13].

**Fig 3.**
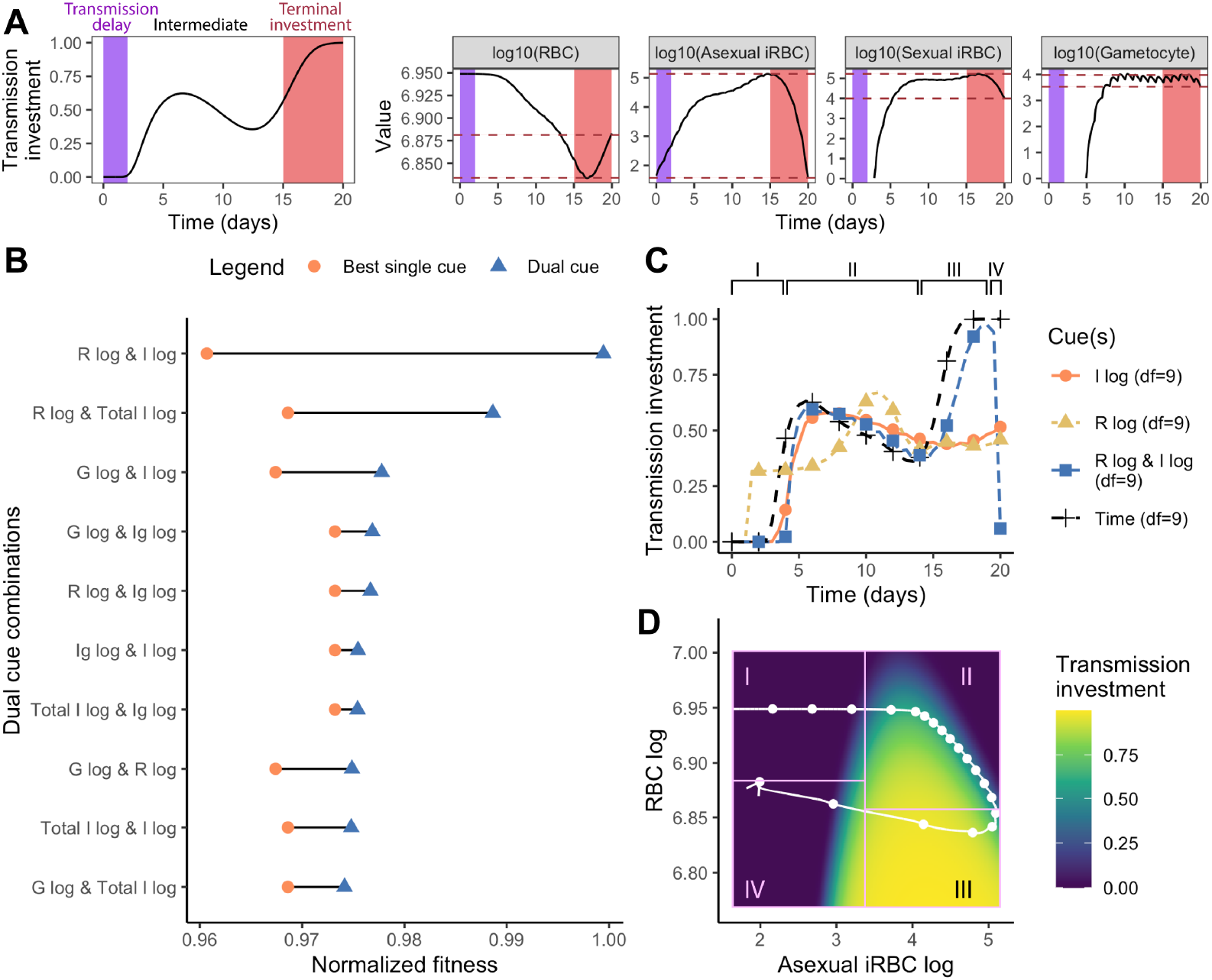
Dual cue perception increases parasite fitness and enables terminal investment. (A) RBC density uniquely marks the onset of terminal investment. In the left panel, we plot transmission investment of parasites sensing time (i.e., the ideal dynamics) and mark its key features: transmission delay (purple) and terminal investment (red). We then plot the dynamics of single cues on the right and mark the range of cue values reached during terminal investment with two red, dashed lines. The overlap of the red dashed lines with earlier cue values indicates that parasites may not be able to uniquely recognize the end of the infection. (B) The fitness of parasites (normalized with respect to the time-varying strategy) adopting the optimal strategy for each dual cue combination is indicated with a blue triangle, while the fitness obtained when perceiving the best performing single cue (of the combination) is represented with an orange circle. To obtain the fitness of a fair comparator time-varying strategy, we optimized transmission investment as a function of time using a spline with nine degrees of freedom (the same degrees of freedom used in the dual cue model), rather than the three degrees of freedom used above and previously [23]. (Note that the marginal fitness gains of using more flexible splines are small [23], but tracking two environmental cues necessitates more degrees of freedom.) (C) The pattern of transmission investment over time for parasites sensing the best dual cue combination (R log & I log, blue square) along with individual components of the cue (I log, orange circle; R log, yellow triangle; each optimized with nine degrees of freedom) and the ideal dynamic (Time, black cross) are plotted together. Parasites sensing the combination of logged asexual iRBC and logged RBC best approximate the time-varying strategy. (D) Transmission investment (colour of heatmap) is plotted as a two-dimensional reaction norm, with logged RBC and logged asexual iRBC as cues. The white line indicates the trajectory of infection, starting at the top left, for parasites adopting this optimal reaction norm, with each point representing a single day. The arrow on the white line indicates the passage of time. For (C-D), the four stages outlined in the results section are annotated with I, II, III, and IV.

To investigate the fitness consequences of sensing two cues, we optimized the transmission investment strategy of parasites sensing all pairwise combinations of log-transformed cues, using two-dimensional spline surfaces and focusing on combinations of log-transformed cues given their higher performance in the single cue optimizations. Our results suggest that sensing dual cue combinations improves parasite fitness relative to sensing a single cue (Fig 3B), with parasites sensing logged RBC and logged iRBC (asexual or total) exhibiting notably higher fitness (Fig 3B). This higher fitness is conferred by transmission investment patterns that are close to the time-varying ideal, with an initial delay in transmission investment and a late infection (nearly) terminal investment before returning to a low transmission investment (Fig 3C, S2 Fig). Increasing the spline flexibility of the single cue models (to match the degrees of freedom in dual cue models) could not reproduce this same pattern (Fig 3C), confirming that dual cue perception specifically enables terminal investment and its associated fitness benefits (S3 Fig).

To better understand the optimal patterns of transmission investment, we next analyzed the reaction norm of the dual cue combination that conferred the highest fitness: logged asexual iRBC and logged RBC. The pattern of transmission investment is defined by a looping journey through the reaction norm surface (Fig 3D), confirming that the cue combination traces a unique path over the course of an infection. We categorized the reaction norm into four distinct sections/stages (Fig 3D). Stage I corresponds to the initial phase of infection, characterized by low transmission investment, low parasite density, and high RBC density (Fig 3C-D). As the infection progresses, the reaction norm enters stage II, characterized by asexual iRBC production and RBC decline. During this stage, the optimal transmission investment increases considerably, followed by subtle declines as the iRBC population continues to increase (Fig 3C-D). The transition from a low to high transmission investment observed as stage I transitions into stage II constitutes delayed transmission investment. Following peak parasite density, the reaction norm enters stage III, which is marked by decreasing iRBC population and gradual replenishment of RBCs, signaling imminent infection clearance. At this stage, transmission investment increases to peak levels (i.e., terminal investment). Finally, as the infection nears clearance and the reaction norm enters stage IV, transmission investment sharply drops as the host heads towards recovery (Fig 3C-D). We note that this drop in predicted transmission investment is likely a consequence of the constraints of the spline surface, and different strategies during this stage would have negligible impact on fitness [23]. Overall, our results suggest that each cue in this combination plays a crucial role in defining progression through infection stages I to III, with asexual iRBC density exhibiting the largest variation in early and late infection (stages I and III), and logged RBC density exhibiting the largest variation during mid-infection (Stage II; Fig 3D). This temporal staggering of iRBC and RBC dynamics (i.e., their hysteretic relationship) ensures that these infection stages can be tracked. As a result, dual cue perception provides parasites with the ability to “state differentiate,” to accurately monitor the progress of the infection and adjust transmission investment accordingly.

Intriguingly, the dual cue combinations that are most informative for parasites making life history decisions (top two in Fig 3B) are also the most informative for predicting patient outcomes in clinical malaria infections [46]. In both cases, two slightly out-of-phase within-host measures can uniquely define a patient’s (or simulated host’s) position on the path from health through sickness to recovery. Plotting this trajectory through disease maps (i.e., the two measures plotted against each other, as in Fig 3D) produces a looping, or hysteretic, pattern, with each infection stage occupying a unique coordinate in this two-dimensional space [46]. To further validate that forming the characteristic looping pattern is a unique feature of our identified best cue combinations, we constructed disease maps using experimental data from laboratory mice infected by a single strain of *P. chabaudi* [11, 42–45, 47] and simulated data from our models where parasite senses time. Both experimental and simulated data indicate that raw or logged RBC with logged iRBC (asexual or total in the model; only total is available in the experimental data) are the only cue combinations that generate fully separated loops (e.g., Fig 4A), while other cue combinations do not (e.g., Fig 4B and C; full data shown in S4 Fig and S5 Fig). When early and late infection stages are “condensed” in space, simulated parasites fail to state differentiate and do not delay transmission investment or terminally invest (S2 Fig). In summary, our results demonstrate that optimal transmission investment requires parasites to unambiguously state differentiate, track infection stages, and dynamically adjust investment in response to changing conditions. Our data suggest that perceiving hysteretic cues — of which a combination of RBC and logged iRBC provides a clean delineation of different infection stages — is a viable method of attaining state differentiation.

**Fig 4.**
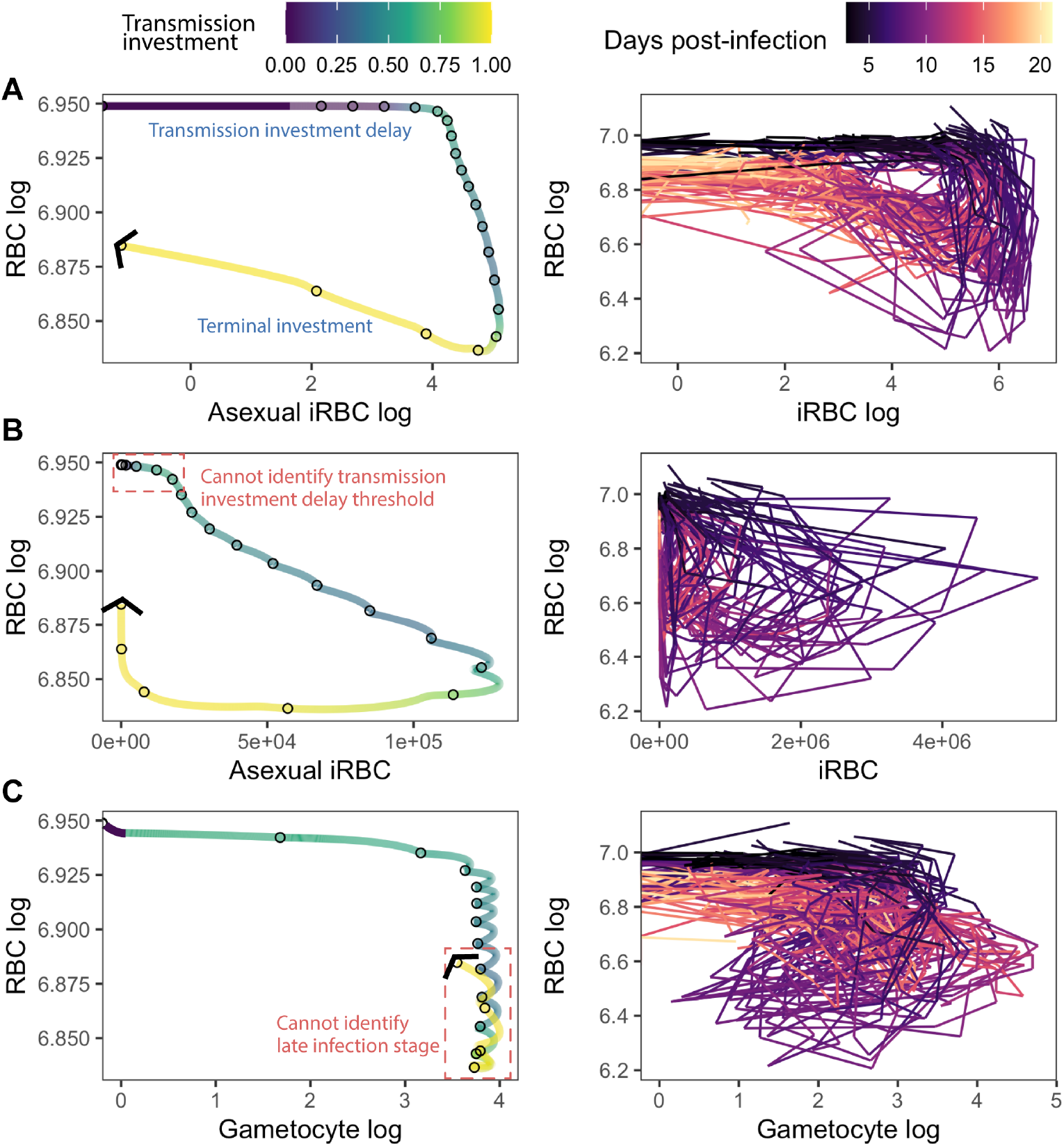
Trajectories of log-transformed RBC and iRBC densities form a looping pattern *in silico* and *in vivo*. Simulated (left) and experimental (right) trajectories of cues that (A) enable transmission delay and terminal investment or (B-C) fail to achieve these patterns. For each panel, the left graph displays the results from our simulations where parasite senses time with nine degrees of freedom, with the distance between each circle representing a single day and the colour of the line representing the observed transmission investment. The right graph displays the infection trajectory of laboratory *P. chabaudi* infections in mice, with the colour of the lines indicating the day post-infection. Note that asexually- and sexually-committed iRBCs were not distinguished in the experimental data. For the simulated trajectories, the first 3 days correspond to the period of delayed transmission investment and the last 5 days correspond to when terminal investment is expected. In (B), early changes in asexual iRBC density cannot be clearly separated (see the red, dotted box), therefore preventing a delayed transmission investment strategy. Similarly, in (C), late state infection stages cannot be separated from earlier time points, preventing parasites from terminally investing.

### Dual cue perception exacerbates the fitness loss incurred by environmental and developmental stochasticity

The benefits of plasticity depend on the quality of information organisms obtain from environmental cues. Adaptive phenotypic plasticity is contingent on proper phenotype–environment matching, where the perceived cue is a reliable predictor of the optimal phenotype [48–52]. Theory suggests that a weak correlation between cue(s) and the optimal phenotype (i.e., low cue reliability) could disfavour plasticity relative to other strategies (i.e., local adaptation [49, 50, 53], canalization [52], bet-hedging [53]). For malaria parasites, there may be unpredictable variation in the environment (e.g., RBC availability [41, 54, 55], immunity [41, 56–58]) or in traits governing parasite development within the host [41]. Such environmental and developmental stochasticity could preclude evolution towards the strategies we predict above. Although our model allows parasites to sense components of their state, and their resulting level of transmission investment will feedback to influence that state, the results above assume that all else is equal between hosts. Specifically, the optimal reaction norms identified above are associated with a particular set of model parameters (the median estimates from posterior distributions in [41]; we refer to this as the deterministic model). This implicitly assumes that hosts are all the same or that parasites are moving between hosts sufficiently quickly that they adapt to an “average” environment.

To investigate how biologically relevant variation affects cue performance, we built a semi-stochastic model based on the work of Kamiya *et al*. [41], which employed a hierarchical Bayesian method to quantify inter-host variation in parasite burst size, erythropoiesis (RBC replenishment), and immune strengths. For all log-transformed single cues and dual cues, we performed 1000 20-day simulations, holding the reaction norm at the optimum from the relevant deterministic simulation, and simultaneously varying the RBC replenishment rate (*ρ*), parasite burst size (*β*), immune activation strengths (*ψ*_*n*_ and *ψ*_*w*_) and immunity half-lives (*ϕ*_*n*_ and *ϕ*_*w*_), sampling parameter values from the reported posterior distributions [41](S6 Fig). Each choice of parameter set will influence the within-host infection dynamics, potentially altering the trajectory of the perceived cue(s), resulting in different, expressed patterns of transmission investment and, thus, fitness. Here we are asking, if parasites are “optimized” to the average host, does environmental and developmental variation change inferences about which cues are best? We quantified the effect of variation on cue performance using the geometric mean fitness, commonly used to assess fitness in fluctuating environments [59, 60].

We find that the geometric mean fitness of parasites responding to a given cue, in the face of this variation, is largely uncoupled from fitness in the deterministic model (i.e., the environment in which the strategy was optimized; Fig 5A), emphasizing that the “best” reaction norm for a certain environment does not always confer the best phenotype in another. Interestingly, all dual cues have a lower geometric mean fitness than the best-performing single cue (I log, Fig 5A). Furthermore, this result emphasizes the challenge of using time as an environmental proxy, since its geometric mean fitness in the semi-stochastic model is among the bottom half of cues. Allowing only one parameter to vary similarly tends to reduce the fitness of the best dual cue strategies disproportionately (S7 Fig). These results indicate a potential tradeoff between fine-tuning fitness in a given environment and susceptibility towards environmental and developmental stochasticity.

**Fig 5.**
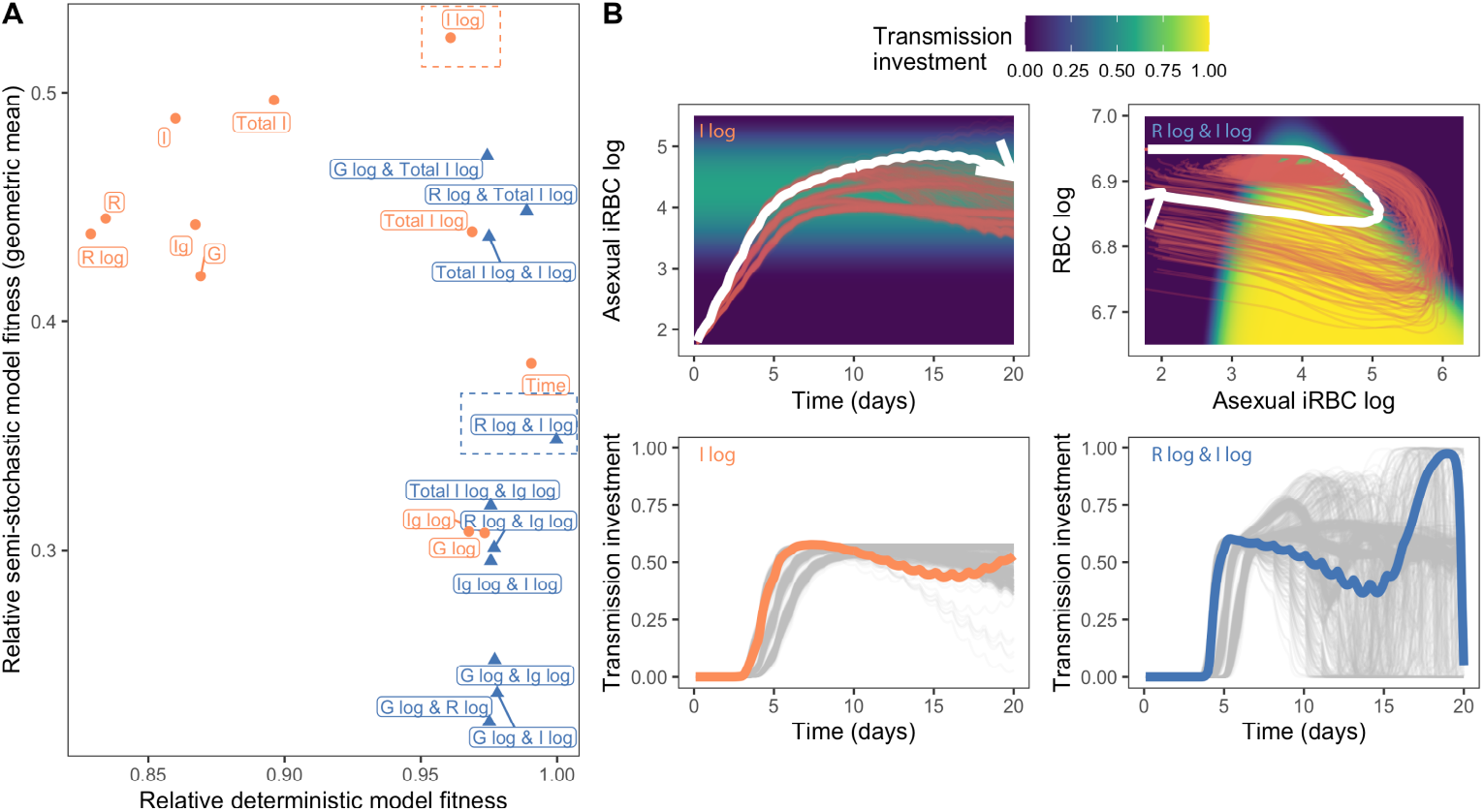
Optimal transmission investment strategies do not always translate effectively across different environmental and developmental conditions. (A) Fitness of parasite strategies in the deterministic model is largely uncoupled from the geometric fitness of those same strategies in the semi-stochastic model. Plotted on the y axis is the geometric mean fitness of 1000 simulations of parasites adopting the optimal transmission investment strategy (sensing time from the deterministic model) using a semi-stochastic model where six parameters are randomly drawn from posterior distributions derived by Kamiya *et al*. [41]. The fitness from the deterministic model is plotted on the x-axis and all values are normalized with respect to the time-varying strategy with nine degrees of freedom (i.e., the dual cue comparator shown in Fig 3C). Note we also plot the fitness of the time-varying strategy with three degrees of freedom. Overall, dual cue-sensing parasites tend to perform worse than single cue-sensing parasites in the face of variation. The cue with the highest geometric fitness (I log) is indicated by a dotted orange box, while the cue combination with the highest deterministic model fitness (R log & I log) is bound by a blue dotted box. (B) Sensing dual cues can increase noise perception and lead to highly variable transmission investment dynamics. The top two graphs display the cue trajectories for parasites sensing the best performing cue in the semi-stochastic simulations (logged asexual iRBC density, plotted against time; left) or dual cues (logged asexual iRBC and logged RBC density, plotted against each other; right). The red lines represent individual semi-stochastic simulation results, while the thicker white line represents the dynamics from the deterministic model. The background colour indicates the optimal reaction norm: on the right, this is the same as what is plotted in Fig 3D. The bottom two graphs display the transmission investment dynamics of the same parasites. The thicker orange/blue line represents the transmission investment trajectory from the deterministic model, while the thinner grey lines represent individual semi-stochastic simulation results.

While sensing two cues provides more information about “where” an infection is, it could also increase the amount of perceived noise and exacerbate the negative effects of environmental/developmental stochasticity. To test whether increased noise perception reduces the fitness of dual cue-sensing parasites, we compared the trajectories of the most “robust” single cue (I log; Fig 5B left panel) and the dual cue combination with the highest deterministic model fitness (R log & I log; Fig 5B right panel). As expected, the variation in asexual iRBC density across our simulations is much lower than that of both asexual iRBC and RBC (Fig 5B top panels). While parasites sensing both types of cues delay transmission, those sensing logged RBC and iRBC display highly variable transmission investment dynamics (Fig 5B bottom right). In contrast, the transmission investment pattern of parasites sensing only logged asexual iRBC resembles that of the deterministic model (Fig 5B bottom left). These results suggest that sensing certain dual cue combinations can reduce parasite fitness by increasing noise perception that induces suboptimal transmission investment.

### Dual cue perception leads to higher virulence

Given that cue perception mediates infection dynamics, it could also influence the severity of infections, or virulence. We estimated virulence using the methods described in Torres *et al*. [46], where infection dynamics are mapped in disease space, with the y-axis quantifying host health (RBC density) and the x-axis quantifying parasite load (total iRBC density). A more virulent infection is expected to produce a higher parasite load while consuming more host resources, tracing a larger loop through disease space. To evaluate the effects of cue perception on virulence, we generated 30-day disease maps of infections, allowing sufficient time for the infection to fully resolve in our model. We use the area enclosed by each disease map as a proxy for virulence, with larger areas indicating higher virulence. We note that we are using “virulence” as synonymous with infection severity and not host mortality, thus we assume no link between virulence and the duration of infection. Recent modeling suggests that resource limitation imposes a more important constraint on the evolution of transmission investment in *P. chabaudi* than does host mortality due to infection [61].

Parasites perceiving only a single cue trace smaller disease maps than parasites that adopt the ideal pattern of transmission investment (by sensing time; Fig 6A). Additionally, the area of the disease map is negatively correlated with parasite fitness across all single cues (Fig 6B), suggesting that if parasites are constrained to perceive a single within-host cue, strategies that result in lower virulence would be favoured. Parasites sensing the top two dual cue combinations produce disease maps that are similar to the map traced by parasites adopting the time-varying strategy (Fig 6C) and result in higher virulence relative to parasites sensing a single cue (Fig 6D; compare x-axis values of points in B and D). Overall, these findings suggest that if parasites have limited information or have evolved to respond to only one cue (e.g., because of physiological constraints or high environmental/developmental variation), virulence will be lower, while if parasites are favoured to more finely track infection stages through dual cue perception, they will exploit host resources to a greater extent and infections will be more virulent.

**Fig 6.**
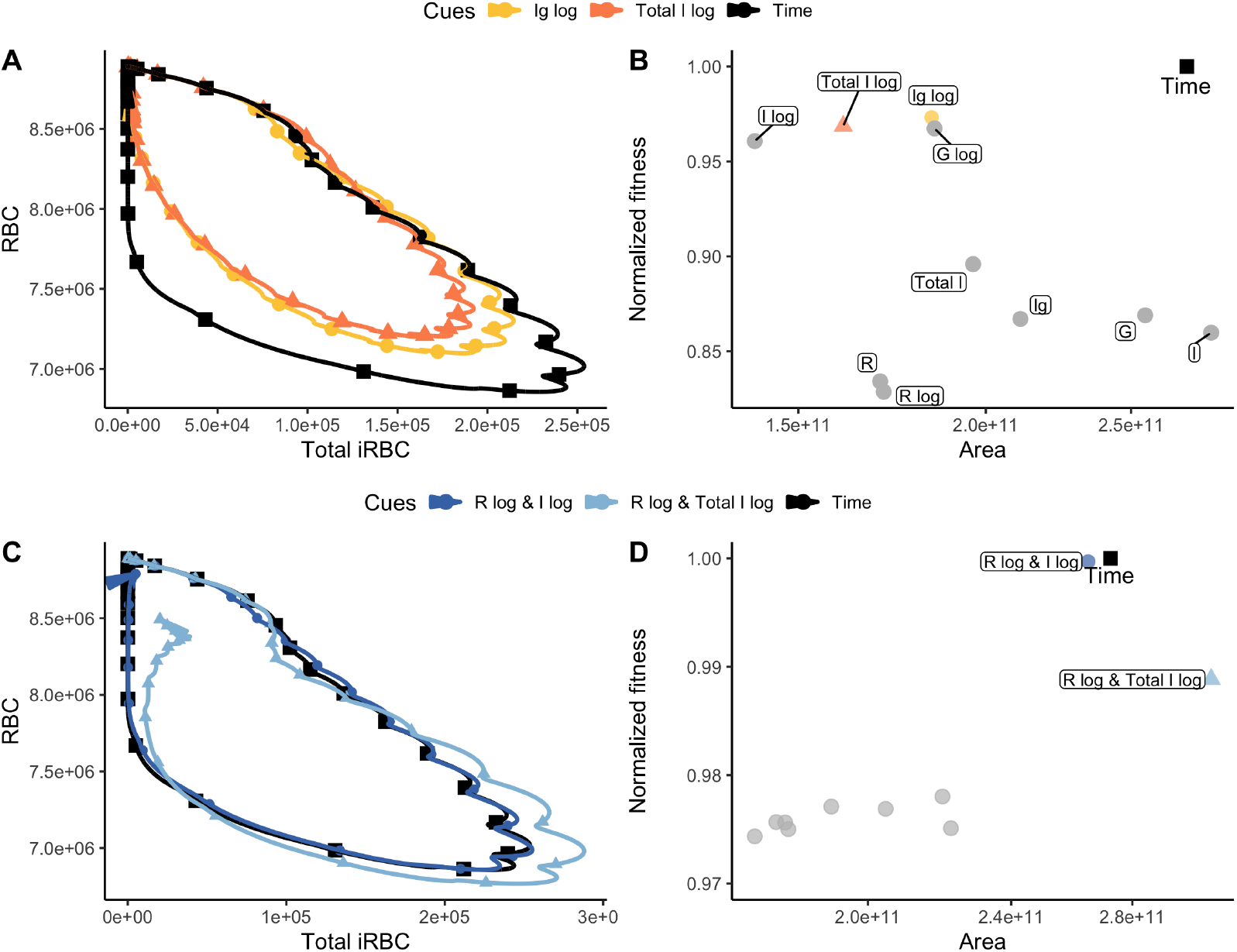
Cue perception mediates infection severity (virulence). (A, C) display the 30-day disease maps of infections where parasites adopt the top two optimal transmission investment strategies when sensing (A) a single cue or (C) dual cues. We use the area of the map traced as a proxy for virulence [46]. The trajectory of parasites that adopt the ideal transmission investment strategy (sensing time) is shown in black. (B, D) Relationship between disease map area and fitness (normalized with respect to the time-varying strategy) for parasites that sense (B) only one cue or (D) dual cues. The top two cues for single and dual cue perceiving parasites are coloured, while the grey points represent the other (B) single or (D) dual cues. Overall, these results suggest that perceiving only a single cue generates selection for strategies that lead to lower virulence while perceiving two cues generates selection for strategies that lead to higher virulence.

## Discussion

Just as with any sexually-reproducing species, investing in reproduction by producing gametocytes is critical for the fitness of *Plasmodium* spp., as this is the only parasite stage capable of being transmitted to mosquito vectors. As for other species, this investment comes with tradeoffs for malaria parasites: a given infected host cell can either produce multiple asexual progeny that can infect new host cells and have the potential to grow or maintain a given infection, or a single, sexually-differentiated gametocyte with the potential to “reproduce” an infection [62]. Thus, maximizing parasite fitness requires balancing these competing demands, the relative importance of which is likely to shift over the course of infection, as the within-host environment changes in response to parasites exploiting the host. Indeed, data show that transmission investment varies throughout infections [11, 12, 26] and appears to change in response to drug treatment in ways predicted to be beneficial for parasites (e.g., [13, 40]). While adaptive plasticity is a well-known solution to the challenge of living in a changing environment (e.g., [63]), there has been skepticism about the fitness benefits of observed parasite plasticity [64], in part because knowledge of parasite environmental sensing mechanisms remains limited. Previous mathematical modeling has treated parasites as either insensitive to environmental change (i.e., constant transmission investment [21, 32–36, 65, 66]) or as perfectly informed (time-varying investment [21, 23, 24]). We instead asked what environmental cues malaria parasites *should* use to set transmission investment, demonstrating that no single cue allows parasites to achieve the best time-varying strategy [23]. Parasites must sense two cues to recapitulate the optimal strategy but that strategy enhances the risk of parasites being “tricked” by environmental and developmental stochasticity into expressing suboptimal phenotypes. We further show that if parasites have the capacity to sense more aspects of their within-host environment, then this has negative consequences for host health.

Previous theoretical work identifies two key features of optimal transmission investment strategies for *P. chabaudi* [23]. First, there should be no investment at the start of infections, when densities are too low to permit successful infection of vectors (*i*.*e*., transmission investment delay). Second, parasites should invest all resources into transmission just before an infection ends, a “terminal investment” strategy from life history theory [7]. Our results delineate the environmental cues that can recapitulate these features. Transmission delay requires a sharp, threshold-dependent transition from no investment to an intermediate investment level as the parasite population grows. Making clear-cut decisions in response to continuous cues is common in biological systems (e.g., cell cycle commitment [67] and developmental patterning [68]) and often relies on ultrasensitivity, characterized by a steep response curve. Our results demonstrate that an ultrasensitive reaction norm where transmission investment rises sharply after a certain population size threshold— a pattern enabled by sensing parasite-centric cues on a logarithmic scale—is sufficient for transmission delay. Prior empirical studies have found that *Plasmodium* transmission investment varies in response to conspecific density [11, 12], parasite-conditioned media [14, 15], and infected red blood cell (iRBC)-derived vesicles [16], suggesting that parasites possess the ability to sense iRBC density. Our results support these responses being part of an adaptive strategy.

In addition to parasite density, *P. chabaudi* has been shown to modulate its transmission investment based on RBC availability and age structure [13, 17–19, 24]. Our results suggest that this is also part of an adaptive strategy, with the combination of iRBC and RBC cues allowing parasites to distinguish between early and late infection. Since most within-host variables go up and down, or vice versa, over time (e.g., iRBC densities rise and then fall due to resource limitation or immunity), one cue cannot provide enough detail to unambiguously establish “where” a parasite is in the course of infection. Combining cues solves this problem. In line with this logic, researchers seeking to define a host’s trajectory over time from health through sickness and on to recovery [69] found a combination of two within-host measures (e.g., parasite density and resource abundance) that are slightly out of phase with each other can uniquely determine a patient’s position on their path for malaria infection [46]. Remarkably, though perhaps predictably, we found that the information desired by medical professionals seeking to predict patient outcomes is also the most informative for parasite life history decisions.

There is a growing appreciation that, across diverse organisms, plasticity is often expressed in response to multiple cues that allow for more finely-tuned strategies [70–72]. Our work adds support to theories suggesting that the adaptive benefits of phenotypic plasticity are enhanced when cues provide non-redundant information [73] and enable organisms to sense distinct states that correspond to specific optimal phenotypes [74]. Yet it is also widely appreciated that integrating multiple cues presents challenges, including the additional metabolic burden of sensing another cue [75] and the multiplicative effect of noise when cues are non-additive [72]. Our complex (decidedly non-additive) dual cue reaction norms present a trade-off: they allow parasites to more accurately recreate optimal strategies in a given environment while rendering them more susceptible to stochastic environmental and developmental change, i.e., host-to-host variability. Such tradeoffs may represent important constraints on the evolution of plastic reproductive investment strategies, perhaps explaining why *P. chabaudi* rarely exhibits *true* terminal investment *in vivo* (displaying only modest increases in transmission investment near infection resolution; [13, 20]), and on the evolution of cue perception more broadly.

The tradeoff hypothesis for virulence evolution stipulates that pathogen transmission inevitably harms the host [76]. While the generality of this framework has been contested (see [77] and references within), empirical data from *Plasmodium* spp. have generally supported a positive relationship between virulence and transmissibility [78–81]. Here, we demonstrate that if parasites are limited to sensing a single cue for setting transmission investment, then the strategies that result in the most onward transmission actually result in less host exploitation. Allowing parasites to sense two cues reverses this relationship and produces the expected pattern of the most onward transmission being achieved by the strategies that also produce the most virulent infections. Importantly, previous work has shown that host heterogeneity can constrain parasite virulence evolution [82–84] and our results suggest another mechanism by which this may be true: responding to two cues—while allowing for state differentiation in one host environment—can dramatically reduce fitness in the face of heterogeneity, potentially favouring single cue (hence, lower virulence) strategies.

Mathematical modeling has enabled us to explore the fitness consequences of cue perception in *P. chabaudi*, and our results suggest where greater model complexity would be most or least informative. For example, incorporating variation in gametocyte circulation time [26] would undercut the reliability of gametocytes as a cue, further supporting our results that it is unlikely to represent the optimal cue. The human malaria parasite *P. falciparum* has a longer sexual development period of 9-12 days [85] and this longer lag between cue perception and phenotype expression will reduce cue reliability [86]. Thus, we anticipate the value of sensing gametocytes to decrease even further in this species. While we have ignored parasite developmental processes that occur in the vector, recent theory suggests they have little qualitative impact on optimal transmission investment strategies [87]. In contrast, the impact of previous exposure on host immunity represents one area where more complexity could qualitatively alter predictions. Our models depict immunologically naive hosts, but where malaria is endemic, parasites will often find themselves in a host with a prior history of infection. Understanding how immune memory influences the quality and reliability of potential within-host cues represents an important area for future research. Our selection of cues assumes that parasites sense only the magnitude of a cue, without detecting temporal changes in that magnitude. Schneider *et al*. [13], in their empirical demonstration of *P. chabaudi* adaptively altering transmission investment, found that replication rate, i.e., the proportional change in parasite densities, had better explanatory power than absolute densities. Broadly speaking, they found transmission investment increased with replication rate, a pattern that is not found with our model since replication rate is highest at the start of infections, before resource or immune limitation kicks in, and not investing in transmission at this point is best. We note that a direct comparison of results is tricky since estimates of transmission investment underlying their observed pattern come from day nine of infections or later. Nonetheless, we identified sensitivity to changes at low population density as a key feature of a “useful” cue and the proportional changes captured by replication rate will similarly share this feature. Another important model assumption is that all parasites perceive the same cue at the same strength, but cues may not be spatially uniform. Variation in signal strength and imperfect cue sensing could lead to heterogeneity in transmission investment in a genotypically uniform parasite population, similar to the dynamics observed in bacterial biofilms [88]. Extrapolating from our results, we hypothesize that if the variation in signal strength is large enough, selection may favour single cue sensing due to the relative robustness to stochastic variation. We note that the variation we incorporated in our semi-stochastic simulations likely underestimates that which would be observed in natural infections, since the dataset fitted by Kamiya *et al*. [41] includes only a single mouse host genetic background studied in a highly controlled experimental system [29]. Thus, environment-phenotype mismatching could be an even larger problem in nature than we have estimated. Additionally, given the variation in burst size and developmental timing across *Plasmodium spp*., there is a great need to quantify the extent of variation across infection parameters to better understand the role of environmental/developmental heterogeneity on parasite evolution.

Beyond malaria, our work presents a generalizable computational approach to study phenotypic plasticity. By defining plastic traits as an optimizable reaction norm, it is possible to explore the fitness consequences of sensing different cues and generate hypotheses regarding why organisms respond to certain cues and not others. Mechanistic models of plasticity that capture feedbacks between traits and the environments that shape them will allow for predicting the consequences of perturbations, which is relevant for understanding not only disease dynamics in the face of control measures, but also the fate of organisms whose environment is rapidly altered by climate change.

## Materials and methods

### Within-host model of *P. chabaudi* infection dynamics

We modified and extended a previously published mathematical model describing the within-host infection dynamics of the rodent malaria parasite, *P. chabaudi* [23] (Fig 1).

The original model consists of ordinary and delay differential equations representing the interactions between the parasite, host red blood cells (RBCs), and host immunity over a 20-day infection. There are three key differences in our model from that previous work. First, we include the activity of two forms of immunity to better capture host defense mechanisms: a targeted immune response that only kills infected RBCs (iRBCs) and an indiscriminate response that kills any RBC, as in [41]. Second, we revised the model to account for the observation that sexually-committed parasites do not engage in gametocytogenesis until the next infection cycle (Next Cycle Conversion; [39]). This necessitated separately tracking the production of sexual merozoites (*M* _g_) and asexual merozoites (*M*) from infected cells and consequently introduced a delay between sexual commitment and the outcome of sexual commitment (i.e., production of an iRBC that will produce a gametocyte, *I*_g_). Third, we modeled transmission investment as a function of one or two within-host cues, instead of time as in [23], to better capture the mechanism of phenotypic plasticity, since there is no evidence malaria parasites track time across multiple generations. We further describe the implications of these changes on the model below.

We depict an immunologically naive host given the availability of experimentally derived parameters. Throughout an infection, we assume indiscriminate immunity eliminates all RBCs (*R, I, I*_*g*_) whereas targeted immunity eliminates only iRBC (*I, I*_*g*_). As in [41], we define *N* and *W* as the proportion of cells cleared per day by indiscriminate and targeted responses, respectively. The magnitude of this activity is dependent on the total abundance of iRBC (*I* + *I*_*g*_):

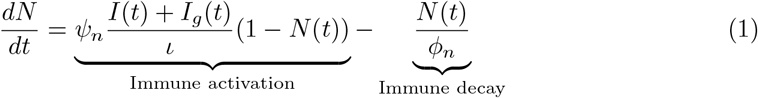

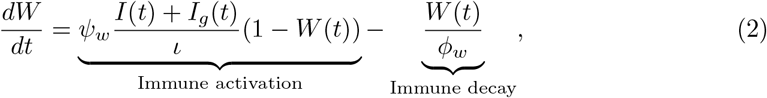

where *ψ*_*n*_ and *ψ*_*w*_ denote the immune activation strengths, *ϕ*_*n*_ and *ϕ*_*w*_ represent the immune activity half-lives, and *ι* represents the theoretical maximum total iRBC density, which determines the threshold at which immunity saturates. Experimental data from mouse infections indicate that indiscriminate immunity is quickly activated but short-lived, whereas targeted immunity has a longer activation period but a longer half-life [41]. Because *N* and *W* have been defined as proportions of cells cleared, they can be rewritten to reflect daily rates of clearance. For example, 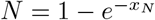, where *x*_*N*_ is equal to the rate of clearance on a given day, and is defined as −ln (1 − *N*), with some rearranging of terms. A similar expression provides the daily clearance rate for targeted killing, *−*ln (1 *− W*).

Mimicking an experimental *P. chabaudi* infection, the host receives an initial dose of asexually-committed iRBC (*I*_0_) to start the infection. We assume no parasites in this initial dose commit to gametocytogenesis and all produce asexual merozoites (*M*). Therefore, before the end of the first asexual development period (*t* ≤*α*, where *α* is the development period of one day for this species), the dynamics of asexual iRBC (*I*) in the host are determined solely by the invasion of RBC by asexual merozoites, natural RBC death, maturation of the initial cohort of infected cells, and immune-mediated killing such that

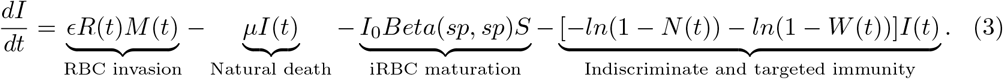

The first term, *ϵR*(*t*)*M* (*t*), represents the production of iRBC through asexual merozoite invasion of RBCs, which occurs at a rate of *ϵ* per day, given contact. Infected RBCs die at a background rate of *µ*, while −*ln*(1 − *N* (*t*)) and −*ln*(1 −*W* (*t*)) describe their rate of death due to indiscriminate and targeted immunity, respectively. The infection-age structure of the initial cohort of asexual iRBCs, denoted as *Beta*(*sp, sp*), is modeled using a Beta distribution with a shape parameter *sp* = 1, allowing iRBC maturation to occur with a uniform probability throughout the development period [89]. The uniform probability of maturation for the initial cohort is computationally efficient and provided extremely similar results to more narrow age distributions in a previous modeling study [23]. The rate of asexual iRBC maturation depends on an exponential survival function *S*, which describes the proportion of iRBC that survives until maturation. *S* is defined by a cumulative hazard function that is calculated by integrating the total rate of death across the development period:

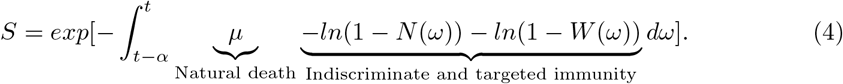

During the period *t ≤ α*, asexual merozoite production is described by

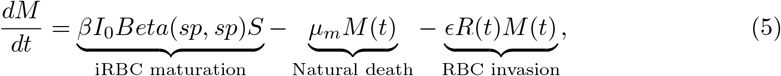

where the first term represents the production of asexually-committed merozoites by asexual iRBC that have survived the duration of the developmental cycle and burst to produce *β* merozoites each. Asexual merozoites are lost by natural death at a rate of *µ*_*m*_ and are removed by their invasion of susceptible RBC.

After all the initially injected iRBC have matured or died (*t > α*), the next cohort of asexual iRBC starts to reach maturity. Transmission investment is defined as the proportion, *c*, of this new generation that commits to gametocytogenesis, resulting in the production of *β* sexually-committed merozoites (*M*_*g*_) per cell. These sexual merozoites invade susceptible RBCs to produce sexual iRBCs (*I*_*g*_). We assume that sexually-committed merozoites have the same death rate (*µ*_*m*_) and rate of RBC invasion (*ϵ*) as asexual merozoites. The remaining proportion (1− *c*) of iRBC produces asexual merozoites and continues the asexual replication cycle. Altogether, when *t > α*, the dynamics of iRBC and merozoites are as follows:

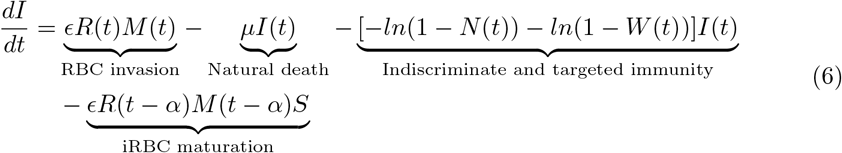

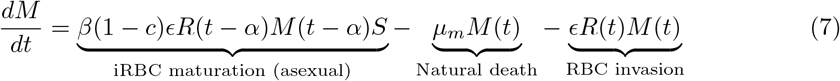

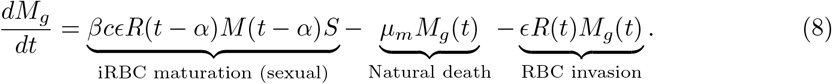

The parameter governing transmission investment (*c*) represents the proportion of iRBCs that produce sexually-committed merozoites and is further defined in the next section.

Before the first cohort of sexual iRBC matures (*α < t* ≤ *α* + *α*_*g*_), the dynamics of sexual iRBC is governed solely by the rate of RBC invasion by sexually-committed merozoites, natural RBC death, and immunity:

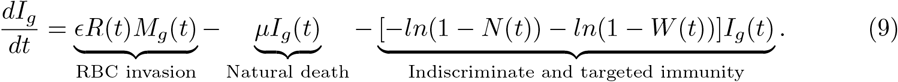

When the first cohort of sexual iRBC matures (*t > α* + *α*_*g*_), each sexual iRBC produces a single gametocyte, *G*, such that

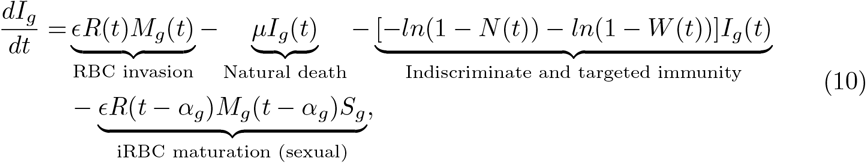

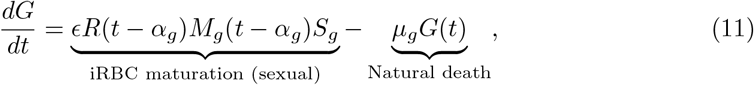

where *µ*_*g*_ is the background death rate of gametocytes. The survival probability of sexual iRBCs during their development cycle is defined by the exponential survival function *S*_*g*_, which is similar to that of asexual iRBC (*S*), except the cumulative hazard function is integrated over the length of sexual iRBC development, *α*_*g*_:

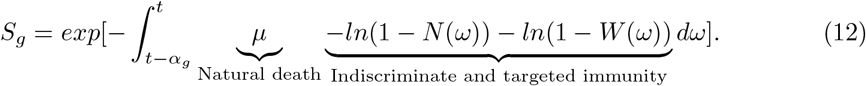

Throughout the infection, the density of uninfected RBCs, *R*, is affected by erythropoiesis, natural death, indiscriminate immunity, and merozoite invasion, such that:

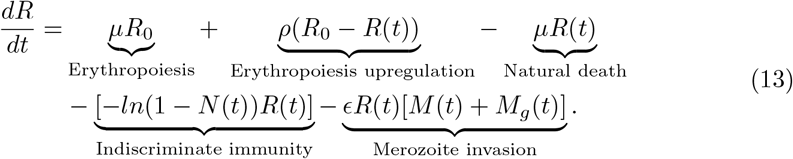

In the absence of parasites, RBCs are replenished at a baseline rate, *µ*, that maintains a homeostatic equilibrium RBC concentration (*R*_0_). During infection, the rate of erythropoiesis increases, and the extent of this upregulation is proportional to the deviation of RBC concentration from equilibrium [90], where *ρ* represents the maximum proportion of the RBC deficit recovered per day. RBCs are eliminated at a natural death rate of *µ* (which balances new production in the absence of infection), removed through indiscriminate immunity (−*ln*(1 −*N* (*t*))), or converted to iRBC via invasion by merozoites.

We parameterized our model based on prior work on malaria intrahost dynamics [23, 41] (Table 2). Because parameter values in Kamiya *et al*. [41] depend on initial dosage (*I*_0_), we estimated our starting dose using the same method, which corresponds to approximately 10^4^-10^5^ iRBCs. Preliminary investigations indicate that varying the initial iRBC dose did not lead to quantitative differences in parasite relative fitness. Consequently, we parameterized *ψ*_*n*_, *ψ*_*w*_, *ϕ*_*n*_, *ϕ*_*w*_ and *ρ* using the median posterior estimates from [41] assuming an initial dosage of 10^4^ iRBCs.

**Table 2.** Parameter values used in the intrahost *P. chabaudi* model.

| Parameter | Definition | Value | Unit | Source |
| --- | --- | --- | --- | --- |
| $R_0$ | Equilibrium RBC concentration | $8.89 \times 10^6$ | RBC/μL | [29] |
| $\mu$ | Background RBC production and RBC/iRBC mortality rate | 0.025 | day<sup>-1</sup> | [57] |
| $p$ | Infection probability of RBC by merozoite invasion | $8 \times 10^{-6}$ | day<sup>-1</sup> | [91] |
| $\alpha$ | Asexual iRBC development period | 1 | day | [92] |
| $\alpha_g$ | Sexual iRBC development period | 2 | day | [93] |
| $\beta$ | Burst size; number of merozoites produced per iRBC | 5.721 | merozoites | [94] |
| $\mu_m$ | Background merozoite mortality rate | 48 | day<sup>-1</sup> | [91] |
| $\mu_g$ | Background gametocyte mortality rate | 4 | day<sup>-1</sup> | [93] |
| $I_0$ | Initial iRBC inoculum size | 43.860 | iRBC/μL | [23] |
| $sp$ | Shape parameter dictating iRBC synchrony | 1 | NA | [89] |
| $\psi_n$ | Activation strength for indiscriminate immunity | 16.692 | NA | [41] |
| $\psi_w$ | Activation strength for targeted immunity | 0.843 | NA | [41] |
| $\phi_n$ | Half-life of indiscriminate immunity | 0.035 | day | [41] |
| $\phi_w$ | Half-life of targeted immunity | 550.842 | day | [41] |
| $\iota$ | Minimum iRBC concentration to induce maximum immunity | $2.18 \times 10^6$ | RBC/μL | [29] |
| $\rho$ | Maximum proportion of RBC recovered per day | 0.263 | NA | [41] |

### Modeling transmission investment

In our model, transmission investment (*c*) is determined by a transformed cue variable, *f*_*t*_, perceived *α* days ago by a parasite infecting a cell, to account for NCC (*f*_*t*_(*t* − *α*)). Following the approach taken by Greischar *et al*. [23], we modeled the transmission investment reaction norm as the complementary log-log basis-spline with three degrees of freedom (*B*_3_). Briefly, this method models the reaction norm as a linear combination of basis function, generating a continuous, smooth cubic curve. The relative contribution of each basis function (also known as the model coefficient) defines the overall shape of the reaction norm (see Figure 1C). We further modify the basis-spline by applying a complementary log-log transformation to constrain the transmission investment value to be between zero and one. This method allows us to use numerical optimization to search for the set of model parameters (i.e., reaction norm shape) that maximizes parasite fitness. Because cubic splines are fundamentally smooth, the resultant reaction norms may have difficulty representing sharp transitions. For that reason, we used untransformed and log-transformed cue values to build our models and capture greater variation in putative reaction norms.

For numerical optimization, the perceived cue must be within the bounds of the reaction norm. To achieve this, the perceived cue variable *f* (*t* − *α*) governing transmission investment is transformed via a linear combination of Heaviside step functions (*H*) from the R package *CRONE* [95] that constrains the cue value to be between zero and the maximum value (max) defined in Table 3. In summary, the transmission investment function is defined as

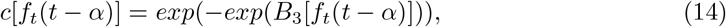

where the Heaviside-transformed environmental cue *f*_*t*_(*t − α*) is given by

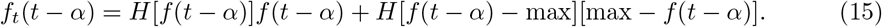

**Table 3.** Within-host environmental cues tested and their range.

| Cue (density per $\mu\text{L}$) | Abbreviation | Min. value | Max. value |
| --- | --- | --- | --- |
| Asexual iRBC | I | 0 | $6 \times 10^6$ |
| Asexual iRBC $\log_{10}$ | I log | 0 | 6.78 |
| Sexual iRBC | Ig | 0 | $6 \times 10^6$ |
| Sexual iRBC $\log_{10}$ | Ig log | 0 | 6.78 |
| Asexual iRBC + sexual iRBC | Total I | 0 | $6 \times 10^6$ |
| Asexual iRBC $\log_{10}$ + sexual iRBC $\log_{10}$ | Total I log | 0 | 6.78 |
| Gametocyte | G | 0 | $6 \times 10^4$ |
| Gametocyte $\log_{10}$ | G log | 0 | 4.78 |
| RBC | R | $10^6$ | $10^7$ |
| RBC $\log_{10}$ | R log | 6 | 7 |

For the best time-varying reaction norm, we defined the perceived cue as the time since infection, which has a range of 0-20 days and a step size of 0.001 (same as the simulation time step). In this case,

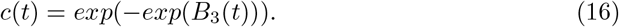

To incorporate dual cue perception, we used the *mgcv* package [96] to model transmission investment as the response variable of a generalized additive model. This model incorporates a full tensor product smooth of two environmental cues with nine degrees of freedom, represented as follows:

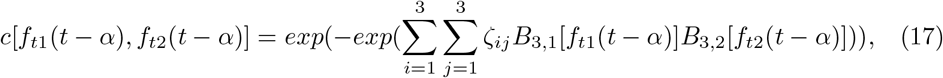

where transmission investment is governed by two Heaviside-transformed cues, *f*_*t*1_ and *f*_*t*2_, perceived *α* days ago. The transmission investment surface (i.e., 2-dimensional reaction norm) is generated by taking the tensor product of the basis-splines of two cues (*B*_3,1_ and *B*_3,2_). The shape of this spline surface is defined by the model coefficients *ζ*_*ij*_.

### Quantifying parasite fitness

To quantify parasite fitness, we calculated the cumulative transmission potential of different transmission investment strategies, following the method described by Greischar *et al*. [23]. For a given time step, transmission potential, *τ* (*t*), is defined as the probability that a mosquito vector becomes infected after biting an infected host and has been shown empirically to be a sigmoid function of gametocyte density in this parasite species [10],

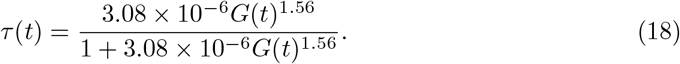

The cumulative transmission potential, or overall fitness of the parasite, is calculated by integrating transmission potential over the entire infection duration of *δ* days:

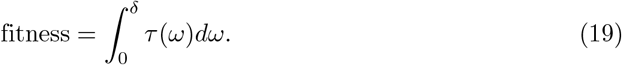

This fitness metric assumes that the mosquito population is at a constant abundance and continuously biting, for the duration of the infection. Although this is a simplification of the evolutionary pressures faced by malaria parasites [35], it enables us to focus our investigation at the within-host level, where transmission investment “decisions” are being made and environmental cues are being perceived by parasites. In our numerical simulations, we estimated fitness with a step size of 0.001 days unless stated otherwise.

### Cue choice and perception

Transmission investment has been shown to vary in response to within-host competition [11, 12] and RBC age structure and availability [13, 17–19, 24], lysophosphatidylcholine level (a key component of the RBC plasma membrane) and exosome-like particles [97, 98]. Whether parasites are responding to these cues directly, or if they are instead a proxy for changes in some other within-host environmental factor is unclear. To capture some of the diversity of putative cues, we defined transmission investment as a function of one of ten different host and parasite-centric measures (Table 3). Additionally, we implemented dual cue models, allowing parasites to simultaneously perceive two different cues. As transmission investment in *P. falciparum* varies non-linearly with cue strength (e.g., *P. falciparum* in response to lysophosphatidylcholine concentration [97]), and within-host dynamics are known to vary only when infectious dose changes by orders of magnitude [29], we introduced non-linear cue perception, enabling parasites to perceive cues either unaltered or log_10_-transformed. We excluded merozoite density as a cue due to its short half-life and the rapid pace of RBC invasion [99]. We also did not include measurements of immunity as cues, given that our model does not track individual components of immunity (rather, it tracks immune killing, phenomenologically), nor do we have sufficient empirical evidence detailing the relationship between different immunity components and transmission investment.

For all cues, we adopt a standardized range that spans the values typically observed when infecting immunologically naive mice with *P. chabaudi* [11], given by the minimum and maximum values in Table 3. All values that exceed the maximum bounds are “perceived” by the parasite as the maximum bound via the modified Heaviside transformation function.

### Fitness optimization

Acute *P. chabaudi* infection tends to resolve within 20 days [100]. For all models, we therefore confined our simulations and optimizations to a 20-day infection period using the parameters detailed in Table 2 and using the *deSolve* package in R [101]. To ensure the convergence and continuity of output dynamics, we simulated the models with a small time step size of 0.001 days using the vode solver [102]. For the dual cue model, we used a larger time step of 0.01 days (which does not greatly alter parasite dynamics but improves computational speed immensely).

To investigate how transmission investment strategies affect parasite fitness, we performed numerical optimization of the basis-spline (single cue) or generalized additive model coefficients (dual cue) to maximize the cumulative transmission potential. We optimized the models using two methods to increase the chances of obtaining the global fitness optima. First, we performed local optimization following the methodology outlined by Greischar *et al*. [23] with the same arbitrary initial values of 0.5 for all spline coefficients, but using the more efficient parallelized L-BFGS-B algorithm (Limited-Memory Broyden–Fletcher–Goldfarb–Shanno algorithm-B) implemented in the *OptimParallel* package [103]. Second, we optimized each model using a hybrid approach to avoid convergence to a local fitness maximum. This technique involved initial global optimization using the Differential Evolution (DE) algorithm from the *DEoptim* package [104] (maximum iterations = 500, step tolerance = 50, differential weighting factor = 0.8, cross-over constant = 0.5 (single cue) or 0.9 (dual cue), population size = 40 (single cue) or 90 (dual cue), time step = 0.01 (single cue) or 0.05 (dual cue)). DE is an efficient and robust algorithm with comparable/better performance relative to other stochastic optimization algorithms [105]. Following optimization by the Differential Evolution algorithm, we further optimized the objective function using the L-BFGS-B algorithm to pinpoint a probable fitness maximum. For the differential Evolution algorithm, we used a parameter bound of (−5, −500, −500, −500) and (5, 500, 5000, 500) for the single cue model and (−10, *−*500, *−*1000, *−*1000, −250, −500, *−*1000, −500, −250) and (10, 500, 1000, 1000, 250, 500, 1000, 500, 250) for the dual cue model. These bounds effectively encompass the majority of basis-spline flexibility. From the two optimization results, we selected the parameter sets that resulted in the highest fitness.

For dual cue models, we implemented an additional optimization method that relies on the single cue models to inform the initial starting point. For dual cue models, we reasoned that the best single cue strategy out of the two provides a theoretical baseline fitness. In other words, the worst a dual-cue strategy should do is as good as the best single cue (out of the two cues), while being unresponsive to the second cue. As such, we applied the L-BFGS-B algorithm to derive a dual cue strategy that “resembles” the best single cue strategy, specifically by minimizing an objective function with two terms: (i) the summed absolute difference between the dual and single cue reaction norms, and (ii) the total standard deviation in transmission investment across the worse cue. Inclusion of the second term ensures that transmission investment is not affected by variations in the worse cue. Together, the objective function is represented by:

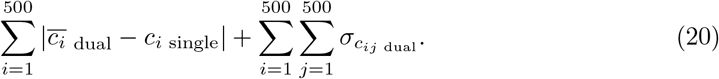

We first discretized the cue ranges into 500 evenly spaced values (*i* = 1…500, better performing cue; *j* = 1…500, worse performing cue). For each cue value with the index *i*, we calculate the deviation of the dual cue model using the absolute difference between 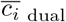, the dual cue transmission investment averaged over the worse cue, and *c*_*i* single_, the corresponding transmission investment in the single cue model. The objective function also quantified the extent to which the worse cue affects transmission investment, specifically by summing the standard deviation across the worse cue value 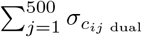 for each cue value indexed *i*.

Following this initial optimization, the resultant parameter is set as the starting point for an additional round of local optimization using the L-BFGS-B algorithm to maximize parasite fitness (same parameters as mentioned above).

As a further validation step, we compared the fitness of each putative single cue optimal strategy to the fitness obtained by 1000 randomly generated strategies. These random strategies were generated within the predefined bounds of (−5, −500, −500, −500) and (5, 500, 5000, 500), established previously to encompass much of the spline space. In no case was the putative best strategy outperformed by a random one (S1 Fig).

### Semi-stochastic infection model

In order to explore the impact of individual infection-level variation on putative optimal transmission investment strategies, we incorporated the following random variables in the within-host model: maximum proportion of RBC deficit recovered per day (*ρ*), parasite burst size (*β*), immunity activation strengths (*ψ*_*n*_ and *ψ*_*w*_), and immunity half-lives (*ϕ*_*n*_ and *ϕ*_*w*_). For each iteration, random variables were drawn from 167 Markov chains (selected at regular intervals from a pool of 16,000 chains) used to estimate the posterior distribution in [41]. For the maximum proportion of RBC recovered per day and parasite burst size, the posterior distributions (originally centered at zero) are re-centered at the deterministic model values of 0.263 and 5.721, respectively. Since the paper from which we draw parameters explored the impact of dose [41], we only considered estimates pertaining to hosts infected with an initial dose of 10^4^ parasites (constituting around 1000 parameter values), which corresponds to our choice for this model parameter (Table 2).

For each optimal transmission investment strategy (i.e., the best strategy for each cue or dual cue combination), we performed 20-day simulations with (i) all parameters allowed to vary from their deterministic default and (ii) only a single parameter varying. Here, we used a time step of 0.01 days, which does not change fitness substantially relative to performing the simulation with a time step of 0.001 days.

### Disease mapping of laboratory mice infections

To explore the dynamics of host and parasite-centric cues in an *in vivo* setting, we curated data from publicly available sources of *P. chabaudi* infection dynamics in mice [11, 42–45, 47]. The curated dataset includes experiments that met the following criteria: a) single parasite strain infection, b) no administration of anti-malarial drugs, and c) availability of data on iRBC, RBC, and gametocyte densities. We then visualized two-dimensional disease maps (e.g., iRBC vs. RBC path plots) using the R package *ggplot2* [106].

### Quantifying parasite virulence

We used the area encompassed within a disease map (where the x-axis displays total iRBC density and the y-axis displays uninfected RBC density, as in [46]) as a proxy for parasite virulence. To calculate this area, we first used the *chull* function [107] to define the convex hull, the smallest set of coordinates that encloses the points within the infection trajectory. We then used the *areapl* function from the *splancs* package [108] to quantify the area of the convex hull.

## Supporting information

**S1 Fig.**
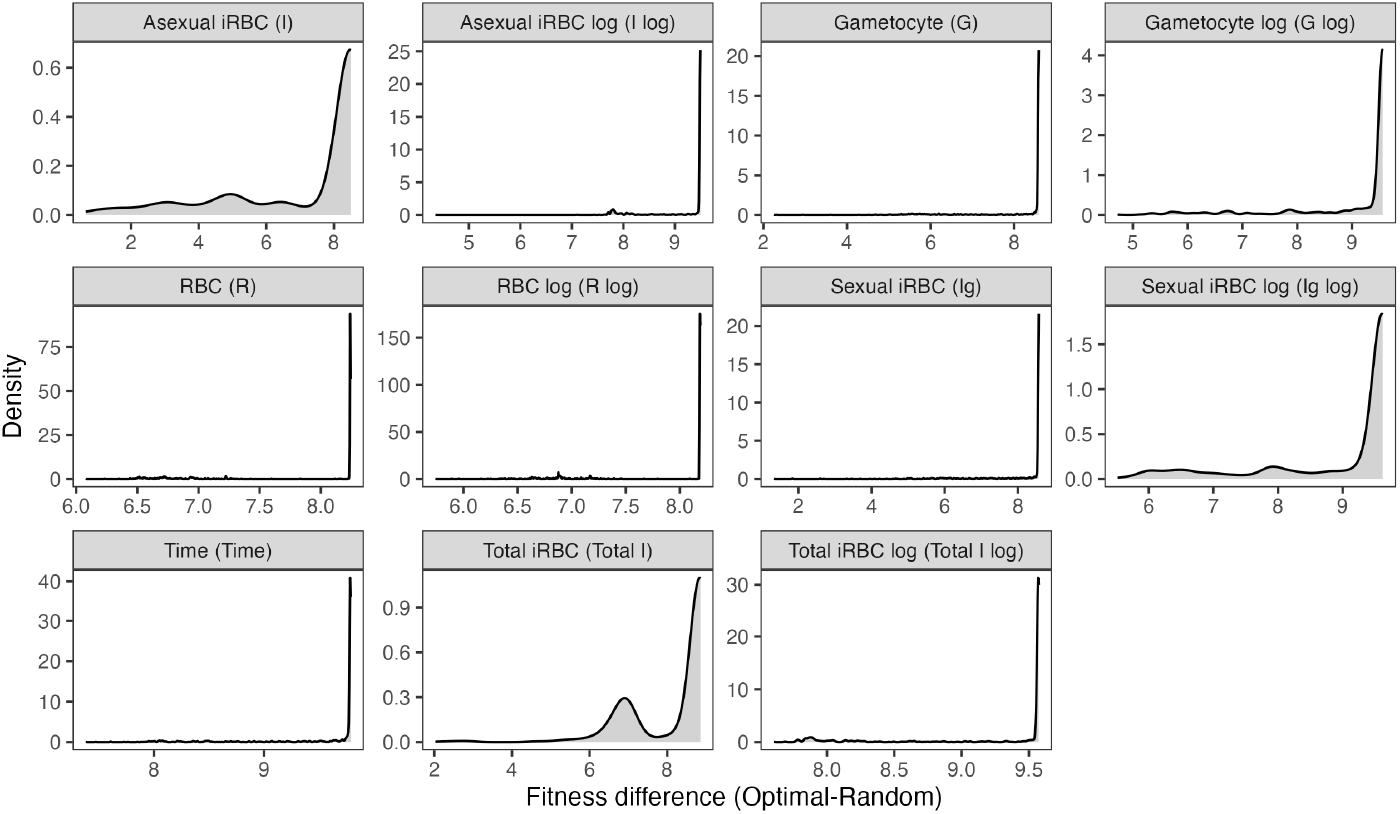
Validation of transmission investment optimization. We conducted 1000 infection simulations for parasites sensing different cues, with each simulation using randomly generated transmission investment strategies. The fitness values of these simulations were compared to the fitness of parasites adopting the optimal transmission investment strategy. Each histogram represents the difference between the optimized fitness and the fitness obtained from the randomly generated strategy. The fitness of the random strategies is always lower.

**S2 Fig.**
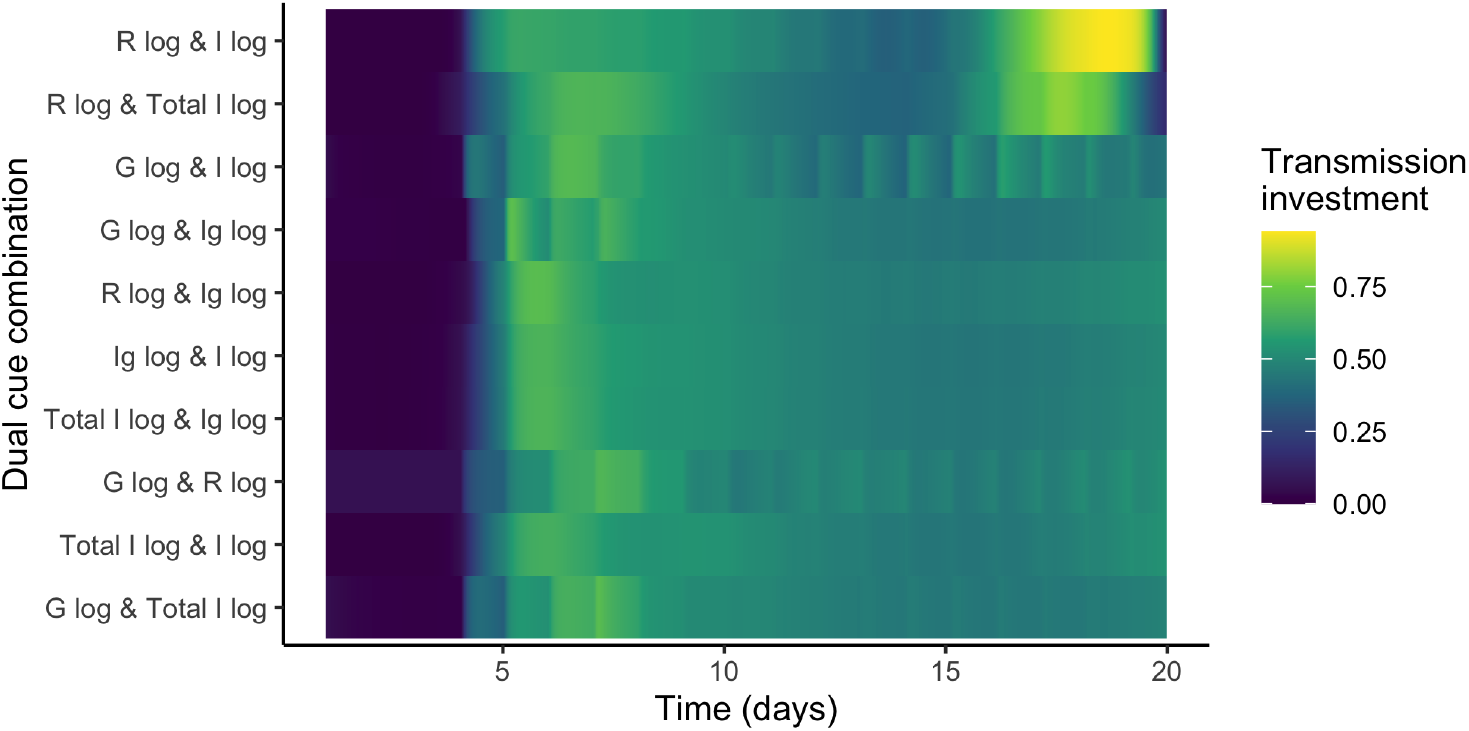
Transmission investment dynamics produced by parasites sensing different dual cue combinations. Transmission investment is plotted for all possible dual cue combinations, with heatmaps ranked in descending order (from top to bottom) based on their associated cumulative transmission potential.

**S3 Fig.**
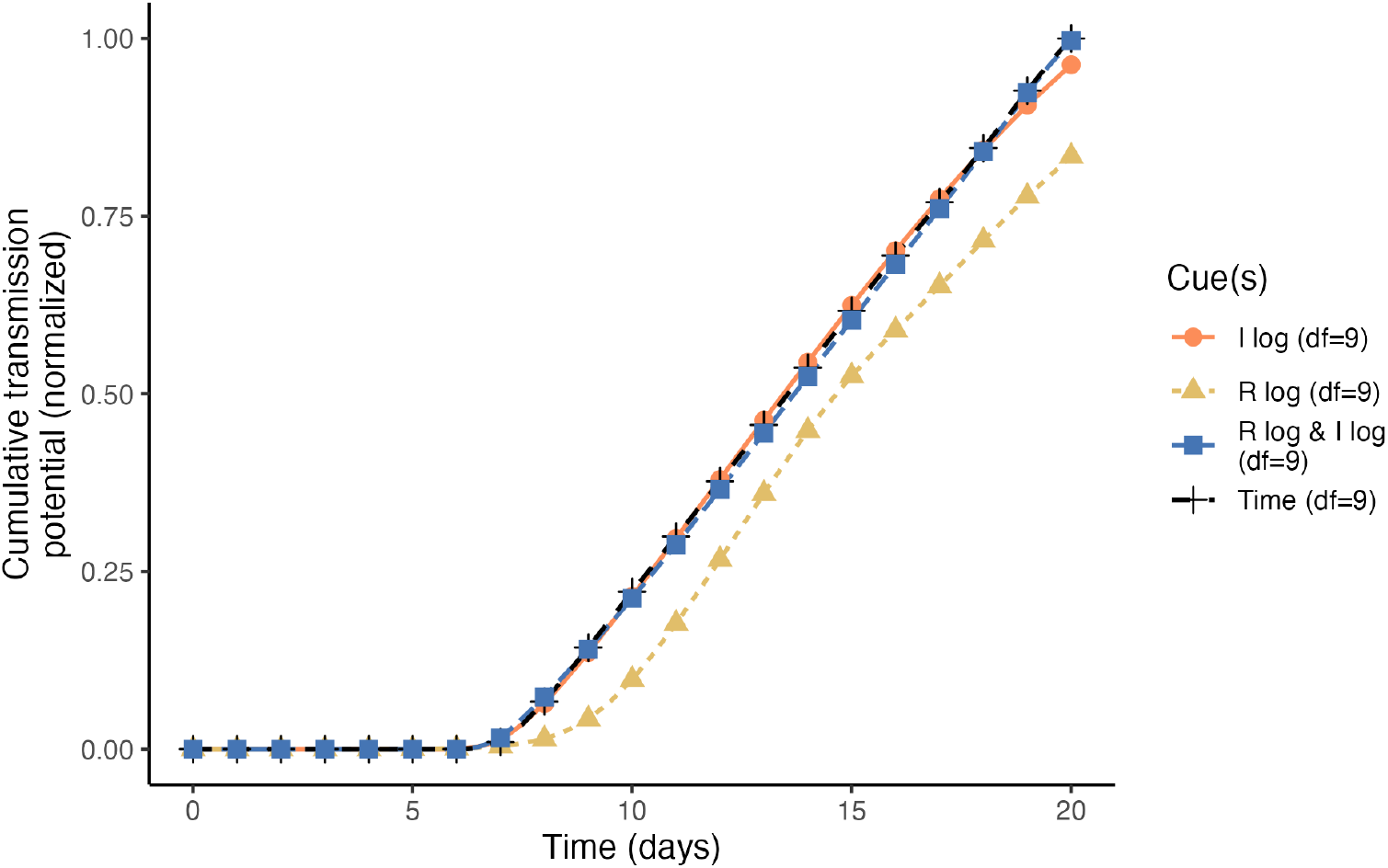
Terminal investment increases the fitness of dual cue-perceiving parasites. The cumulative transmission potential of parasites sensing time, the best performing dual cue combination (R log & I log), or the individual single cues (I log, R log) are indicated with a black cross, blue square, orange circle, and yellow triangle, respectively. The cumulative transmission potential is normalized with respect to the fitness of the best time-varying strategy. Note that time, the dual cue combination, and I log all permit an initial delay in transmission investment, while R log does not; the fitness advantage of this delayed strategy can be clearly viewed on these plots as fitness accrues faster. Only time and the dual cue combination lead to terminal investment, leading to additional fitness gains from day 18 onward.

**S4 Fig.**
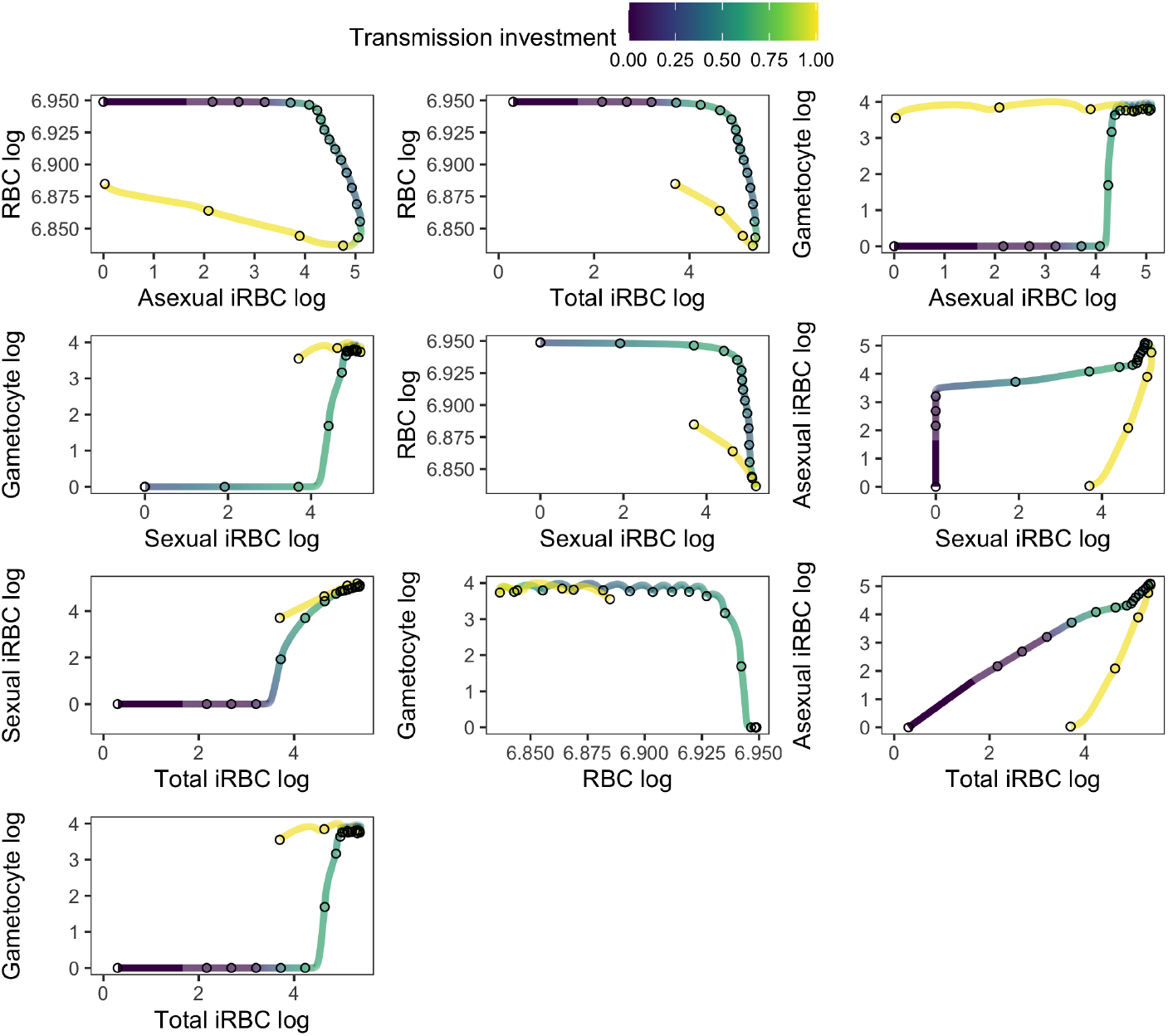
Logged RBC and logged iRBC form a looping trajectory *in silico*. Graphs display the two-dimensional reaction norm and infection trajectories for parasite sensing time with nine degrees of freedom. The trajectories depict the dynamics of the cues (indicated on the x- and y-axes) along with the corresponding transmission investment (represented by the colour of the line). Note that the dynamics of gametocytes and sexual iRBC are hard to display completely given that these densities start off at zero. The distance between each hollow circle represents one day.

**S5 Fig.**
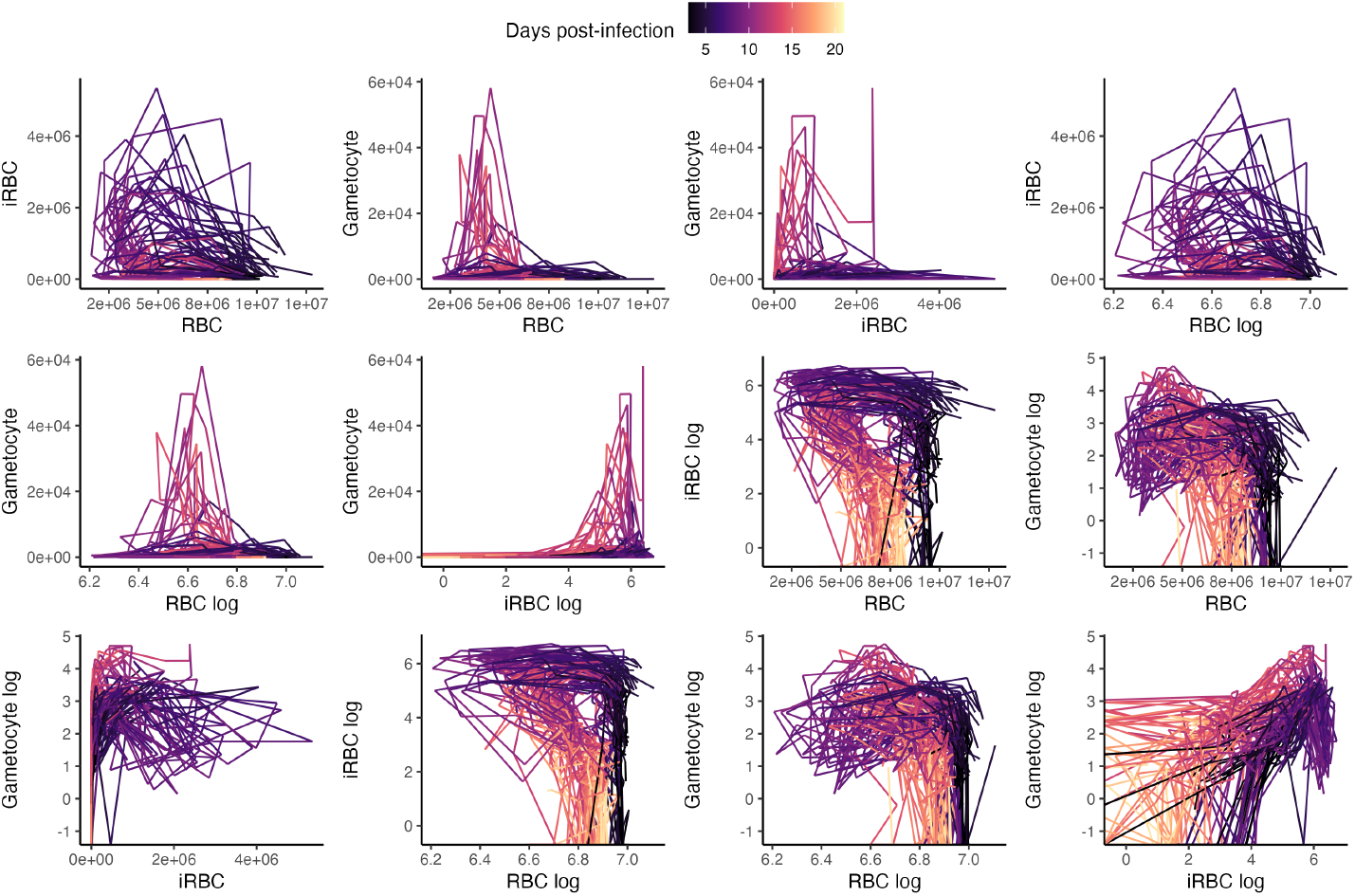
RBC/logged RBC and logged iRBC form a looping trajectory *in vivo*. To identify combinations of within-host variables that exhibit a looping relationship, we plotted all possible permutations of experimentally derived *P. chabaudi* variables. Each line represents the dynamics of a single strain of infecting parasite. The line colour indicates the progression of time.

**S6 Fig.**
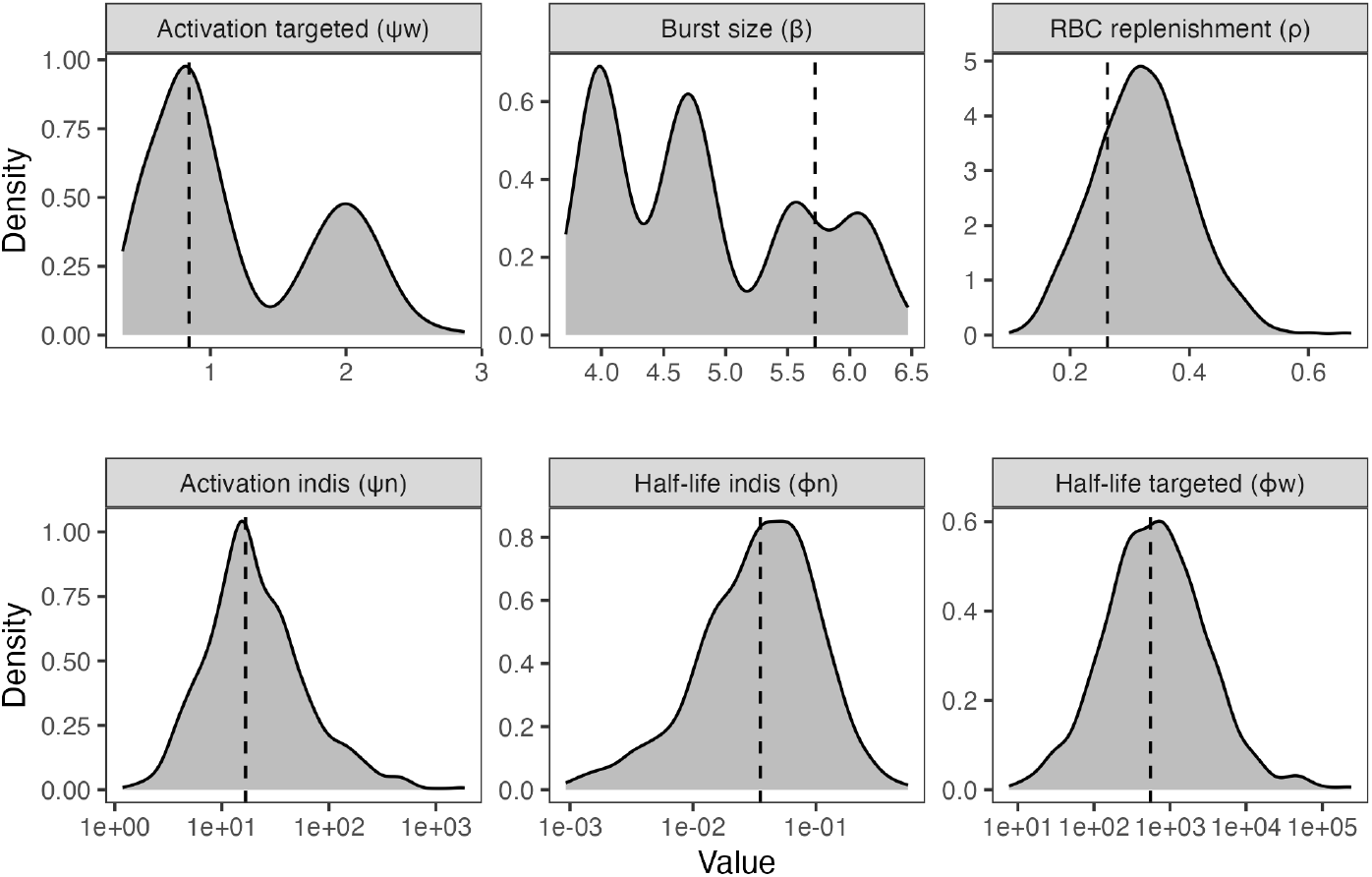
Posterior distribution of parameter values used for the semi-stochastic model. The histograms display the distribution of 1000 parameter values drawn from 167 Markov chains derived by Kamiya *et al* [41]. The parameter values used in the deterministic model are indicated by the dotted line.

**S7 Fig.**
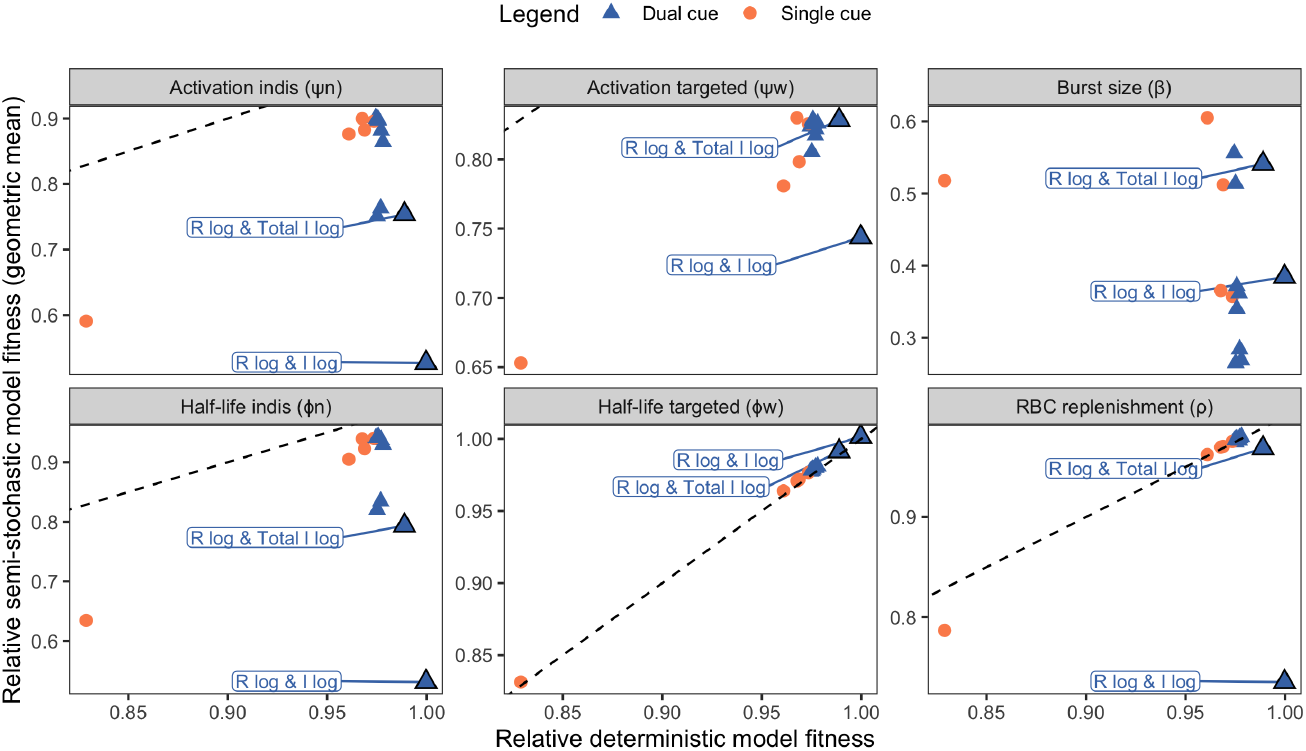
Perceiving the optimal dual cue combinations increases parasite susceptibility to environmental and developmental stochasticity. We conducted 1000 simulations, each lasting 20 days, in which parasites adopted the optimal transmission investment strategy (from the deterministic model) using a semi-stochastic model where only one parameter was randomly drawn from the posterior distributions (see S6 Fig). For each parameter that is varied (indicated in the facet panels), we plot the geometric mean of the fitness on the y-axis and the fitness from the deterministic model on the x-axis. All fitness values are normalized with respect to the best time-varying strategy. The dashed line represents a 1:1 relationship between the x-axis and the y-axis; the line is not visible on the plot for burst size, since variation in this parameter has the greatest negative influence on fitness.

## Acknowledgments

We thank members of the Mideo Lab for useful feedback on the project.

## References

1. Jones OR, Scheuerlein A, Salguero-Gómez R, Camarda CG, Schaible R, Casper BB, et al. Diversity of ageing across the tree of life. Nature. 2014;505(7482):169–173. doi:10.1038/nature12789.

2. Gadgil M, Bossert WH. Life Historical Consequences of Natural Selection. The American Naturalist. 1970;104(935):1–24. doi:10.1086/282637.

3. Clutton-Brock TH, Albon SD, Guinness FE. Maternal dominance, breeding success and birth sex ratios in red deer. 1984;308(5957):358–360. doi:10.1038/308358a0.

4. McNamara JM, Houston AI. State-dependent life histories. Nature. 1996;380(6571):215–221. doi:10.1038/380215a0.

5. Cotter SC, Ward RJS, Kilner RM. Age-specific reproductive investment in female burying beetles: independent effects of state and risk of death. Functional Ecology. 2011;25(3):652–660. doi:10.1111/j.1365-2435.2010.01819.x.

6. Velando A, Drummond H, Torres R. Senescent birds redouble reproductive effort when ill: confirmation of the terminal investment hypothesis. Proceedings of the Royal Society B: Biological Sciences. 2006;273(1593):1443–1448. doi:10.1098/rspb.2006.3480.

7. Carter LM, Kafsack BFC, Llinás M, Mideo N, Pollitt LC, Reece SE. Stress and sex in malaria parasites: Why does commitment vary? Evolution, Medicine, and Public Health. 2013;2013(1):135–147. doi:10.1093/EMPH/EOT011.

8. Churcher TS, Bousema T, Walker M, Drakeley C, Schneider P, Ouédraogo AL, et al. Predicting mosquito infection from Plasmodium falciparum gametocyte density and estimating the reservoir of infection. eLife. 2013;2:e00626. doi:10.7554/eLife.00626.

9. Bell AS, Huijben S, Paaijmans KP, Sim DG, Chan BHK, Nelson WA, et al. Enhanced Transmission of Drug-Resistant Parasites to Mosquitoes following Drug Treatment in Rodent Malaria. PLOS ONE. 2012;7(6):e37172. doi:10.1371/journal.pone.0037172.

10. Huijben S, Nelson WA, Wargo AR, Sim DG, Drew DR, Read AF. Chemotherapy, within-host ecology and the fitness of drug-resistant malaria parasites. Evolution. 2010;64(10):2952–2968. doi:10.1111/J.1558-5646.2010.01068.X.

11. Pollitt LC, Mideo N, Drew DR, Schneider P, Colegrave N, Reece SE. Competition and the Evolution of Reproductive Restraint in Malaria Parasites. The American Naturalist. 2011;177(3):358–367. doi:10.1086/658175.

12. Cameron A, Reece SE, Drew DR, Haydon DT, Yates AJ. Plasticity in transmission strategies of the malaria parasite, Plasmodium chabaudi : environmental and genetic effects. Evolutionary Applications. 2013;6(2):365. doi:10.1111/EVA.12005.

13. Schneider P, Greischar MA, Birget PLG, Repton C, Mideo N, Reece SE. Adaptive plasticity in the gametocyte conversion rate of malaria parasites. PLOS Pathogens. 2018;14(11):e1007371. doi:10.1371/JOURNAL.PPAT.1007371.

14. Williams JL. Stimulation of Plasmodium falciparum gametocytogenesis by conditioned medium from parasite cultures. The American Journal of Tropical Medicine and Hygiene. 1999;60(1):7–13. doi:10.4269/ajtmh.1999.60.7.

15. Dyer M, Day Karen P. Regulation of the rate of asexual growth and commitment to sexual development by diffusible factors from in vitro cultures of Plasmodium falciparum. The American Journal of Tropical Medicine and Hygiene. 2003;68(4):403–409. doi:10.4269/AJTMH.2003.68.403.

16. Mantel PY, Hoang A, Goldowitz I, Potashnikova D, Hamza B, Vorobjev I, et al. Malaria-Infected Erythrocyte-Derived Microvesicles Mediate Cellular Communication within the Parasite Population and with the Host Immune System. Cell Host & Microbe. 2013;13(5):521–534. doi:10.1016/J.CHOM.2013.04.009.

17. Trager W, Gill GS. Enhanced Gametocyte Formation in Young Erythrocytes by Plasmodium falciparum In Vitro. The Journal of Protozoology. 1992;39(3):429–432. doi:10.1111/j.1550-7408.1992.tb01476.x.

18. Trager W, Gill GS, Lawrence C, Nagel RL. Plasmodium falciparum: Enhanced Gametocyte Formationin Vitroin Reticulocyte-Rich Blood. Experimental Parasitology. 1999;91(2):115–118. doi:10.1006/expr.1998.4347.

19. Reece SE, Duncan AB, West SA, Read AF. Host cell preference and variable transmission strategies in malaria parasites. Proceedings of the Royal Society B: Biological Sciences. 2005;272(1562):511–517. doi:10.1098/rspb.2004.2972.

20. Buckling A, Crooks L, Read A. Plasmodium chabaudi: Effect of Antimalarial Drugs on Gametocytogenesis. Experimental Parasitology. 1999;93(1):45–54. doi:10.1006/expr.1999.4429.

21. Koella JC, Antia R. Optimal Pattern of Replication and Transmission for Parasites with Two Stages in Their Life Cycle. Theoretical Population Biology. 1995;47(3):277–291. doi:10.1006/tpbi.1995.1012.

22. Reece SE, Ramiro RS, Nussey DH. Synthesis: Plastic parasites: sophisticated strategies for survival and reproduction? Evolutionary Applications. 2009;2(1):11–23. doi:10.1111/j.1752-4571.2008.00060.x.

23. Greischar MA, Mideo N, Read AF, Bjørnstad ON. Predicting optimal transmission investment in malaria parasites. Evolution. 2016;70(7):1542–1558. doi:10.1111/EVO.12969.

24. Birget PLG, Repton C, O’Donnell AJ, Schneider P, Reece SE. Phenotypic plasticity in reproductive effort: malaria parasites respond to resource availability. Proceedings of the Royal Society B: Biological Sciences. 2017;284(1860). doi:10.1098/RSPB.2017.1229.

25. Clutton-Brock TH. Reproductive Effort and Terminal Investment in Iteroparous Animals. https://doiorg/101086/284198. 2015;123(2):p212–229. doi:10.1086/284198.

26. Eichner M, Diebner HH, Molineaux L, Collins WE, Jeffery GM, Dietz K. Genesis, sequestration and survival of Plasmodium falciparum gametocytes: parameter estimates from fitting a model to malariatherapy data. Transactions of The Royal Society of Tropical Medicine and Hygiene. 2001;95(5):497–501. doi:10.1016/S0035-9203(01)90016-1.

27. Greischar MA, Mideo N, Read AF, Bjørnstad ON. Quantifying Transmission Investment in Malaria Parasites. PLOS Computational Biology. 2016;12(2):e1004718. doi:10.1371/JOURNAL.PCBI.1004718.

28. Pianka ER, Parker WS. Age-Specific Reproductive Tactics. The American Naturalist. 1975;109(968):453–464. doi:10.1086/283013.

29. Timms R, Colegrave N, Chan BHK, Read AF. The effect of parasite dose on disease severity in the rodent malaria Plasmodium chabaudi. Parasitology. 2001;123(Pt 1):1–11. doi:10.1017/S0031182001008083.

30. Metcalf CJE, Graham AL, Huijben S, Barclay VC, Long GH, Grenfell BT, et al. Partitioning regulatory mechanisms of within-host malaria dynamics using the effective propagation number. Science. 2011;333(6045):984–988. doi:10.1126/SCIENCE.1204588.

31. Barclay VC, Råberg L, Chan BHK, Brown S, Gray D, Read AF. CD4+T cells do not mediate within-host competition between genetically diverse malaria parasites. Proceedings of the Royal Society B: Biological Sciences. 2008;275(1639):1171–1179. doi:10.1098/rspb.2007.1713.

32. McKenzie FE, Bossert WH. An integrated model of Plasmodium falciparum dynamics. Journal of Theoretical Biology. 2005;232(3):411–426. doi:10.1016/j.jtbi.2004.08.021.

33. Bousema T, Drakeley C. Epidemiology and infectivity of Plasmodium falciparum and Plasmodium vivax gametocytes in relation to malaria control and elimination. Clinical Microbiology Reviews. 2011;24(2):377–410. doi:10.1128/CMR.00051-10.

34. Cao P, Collins KA, Zaloumis S, Wattanakul T, Tarning J, Simpson JA, et al. Modeling the dynamics of Plasmodium falciparum gametocytes in humans during malaria infection. eLife. 2019;8:e49058. doi:10.7554/eLife.49058.

35. Greischar MA, Beck-Johnson LM, Mideo N. Partitioning the influence of ecology across scales on parasite evolution. Evolution. 2019;73(11):2175–2188. doi:10.1111/EVO.13840.

36. Early AM, Camponovo F, Pelleau S, Cerqueira GC, Lazrek Y, Volney B, et al. Declines in prevalence alter the optimal level of sexual investment for the malaria parasite Plasmodium falciparum. Proceedings of the National Academy of Sciences. 2022;119(30):e2122165119. doi:10.1073/pnas.2122165119.

37. Lensen A, Bril A, van de Vegte M, van Gemert GJ, Eling W, Sauerwein R. Plasmodium falciparum: infectivity of cultured, synchronized gametocytes to mosquitoes. Experimental Parasitology. 1999;91(1):101–103. doi:10.1006/expr.1998.4354.

38. de Boor C. A Practical Guide to Splines. 1st ed. No. 27 in Applied Mathematical Sciences. Springer New York; 2001.

39. Dixon MWA, Thompson J, Gardiner DL, Trenholme KR. Sex in Plasmodium: a sign of commitment. Trends in Parasitology. 2008;24(4):168–175. doi:10.1016/J.PT.2008.01.004.

40. Birget PLG, Greischar MA, Reece SE, Mideo N. Altered life history strategies protect malaria parasites against drugs. Evolutionary Applications. 2018;11(4). doi:10.1111/eva.12516.

41. Kamiya T, Greischar MA, Schneider DS, Mideo N. Uncovering drivers of dose-dependence and individual variation in malaria infection outcomes. PLOS Computational Biology. 2020;16(10):e1008211. doi:10.1371/JOURNAL.PCBI.1008211.

42. Huijben S, Sim DG, Nelson WA, Read AF. Data from: The fitness of drug-resistant malaria parasites in a rodent model: multiplicity of infection. Dryad. 2011;.

43. Huijben S, Nelson WA, Wargo AR, Sim DG, Drew DR, Read AF. Data from: Chemotherapy, within-host ecology and the fitness of drug-resistant malaria parasites. Dryad. 2013;.

44. Huijben S, Bell AS, Sim DG, Tomasello D, Mideo N, Day T, et al. Data from: Aggressive chemotherapy and the selection of drug resistant pathogens. Dryad. 2014;.

45. Huijben S, Chan BHK, Nelson WA, Read AF. Data from: The impact of within-host ecology on the fitness of a drug-resistant parasite. Dryad. 2018;.

46. Torres BY, Oliveira JHM, Thomas Tate A, Rath P, Cumnock K, Schneider DS. Tracking Resilience to Infections by Mapping Disease Space. PLOS Biology. 2016;14(4):e1002436. doi:10.1371/JOURNAL.PBIO.1002436.

47. Reece SE, Drew DR, Gardner A. Sex ratio adjustment and kin discrimination in malaria parasites. Nature. 2008;453(7195). doi:10.1038/nature06954.

48. Moran NA. The Evolutionary Maintenance of Alternative Phenotypes. The American Naturalist. 1992;139(5):971–989. doi:10.1086/285369.

49. Scheiner SM. The genetics of phenotypic plasticity. VII. Evolution in a spatially-structured environment. Journal of Evolutionary Biology. 1998;11(3):303–320. doi:10.1046/j.1420-9101.1998.11030303.x.

50. De Jong G. Unpredictable selection in a structured population leads to local genetic differentiation in evolved reaction norms. Journal of Evolutionary Biology. 1999;12(5). doi:10.1046/j.1420-9101.1999.00118.x.

51. Schlaepfer MA, Runge MC, Sherman PW. Ecological and evolutionary traps. Trends in Ecology & Evolution. 2002;17(10):474–480. doi:10.1016/S0169-5347(02)02580-6.

52. Reed TE, Robin SW, Schindler DE, Hard JJ, Kinnison MT. Phenotypic plasticity and population viability: The importance of environmental predictability. Proceedings of the Royal Society B: Biological Sciences. 2010;277(1699). doi:10.1098/rspb.2010.0771.

53. Tufto J. The evolution of plasticity and nonplastic spatial and temporal adaptations in the presence of imperfect environmental cues. American Naturalist. 2000;156(2). doi:10.1086/303381.

54. Antia R, Yates A, de Roode JC. The dynamics of acute malaria infections. I. Effect of the parasite’s red blood cell preference. Proceedings of the Royal Society B: Biological Sciences. 2008;275(1641):1449–1458. doi:10.1098/rspb.2008.0198.

55. Birget PLG, Kimberley F Prior, Savill NJ, Steer L, Reece SE. Plasticity and genetic variation in traits underpinning asexual replication of the rodent malaria parasite, Plasmodium chabaudi. Malaria Journal. 2019;18(1):222. doi:10.1186/s12936-019-2857-0.

56. Haydon DT, Matthews L, Timms R, Colegrave N. Top-down or bottom-up regulation of intra-host blood-stage malaria: do malaria parasites most resemble the dynamics of prey or predator? Proceedings of the Royal Society B: Biological Sciences. 2003;270(1512):289–298. doi:10.1098/rspb.2002.2203.

57. Miller MR, Råberg L, Read AF, Savill NJ. Quantitative Analysis of Immune Response and Erythropoiesis during Rodent Malarial Infection. PLOS Computational Biology. 2010;6(9):e1000946. doi:10.1371/JOURNAL.PCBI.1000946.

58. Kamiya T, Davis NM, Greischar MA, Schneider D, Mideo N. Linking functional and molecular mechanisms of host resilience to malaria infection. eLife. 2021;10. doi:10.7554/ELIFE.65846.

59. Lewontin RC, Cohen D. On population growth in a randomly varying environment. Proceedings of the National Academy of Sciences of the United States of America. 1969;62(4):1056–1060. doi:10.1073/pnas.62.4.1056.

60. Gillespie J. Polymorphism in random environments. Theoretical Population Biology. 1973;4(2):193–195. doi:10.1016/0040-5809(73)90028-2.

61. Pak D, Kamiya T, Greischar MA. Proliferation in malaria parasites: How resource limitation can prevent evolution of greater virulence. Evolution. 2024;78(7):1287–1301. doi:10.1093/evolut/qpae057.

62. Schneider P, Reece SE. The private life of malaria parasites: Strategies for sexual reproduction. Molecular and Biochemical Parasitology. 2021;244:111375. doi:10.1016/J.MOLBIOPARA.2021.111375.

63. Pigliucci M. Evolution of phenotypic plasticity: where are we going now? Trends in Ecology & Evolution. 2005;20(9):481–486. doi:10.1016/j.tree.2005.06.001.

64. Kochin BF, Bull JJ, Antia R. Parasite Evolution and Life History Theory. PLOS Biology. 2010;8(10):e1000524. doi:10.1371/journal.pbio.1000524.

65. McKenzie FE, Bossert WH. The Optimal Production of Gametocytes by Plasmodium falciparum. Journal of Theoretical Biology. 1998;193(3):419–428. doi:10.1006/jtbi.1998.0710.

66. Mideo N, Day T. On the evolution of reproductive restraint in malaria. Proceedings of the Royal Society B: Biological Sciences. 2008;275(1639):1217–1224. doi:10.1098/RSPB.2007.1545.

67. Charvin G, Oikonomou C, Siggia ED, Cross FR. Origin of Irreversibility of Cell Cycle Start in Budding Yeast. PLOS Biology. 2010;8(1):e1000284. doi:10.1371/journal.pbio.1000284.

68. Melen GJ, Levy S, Barkai N, Shilo B. Threshold responses to morphogen gradients by zero-order ultrasensitivity. Molecular Systems Biology. 2005;1(1):2005.0028. doi:10.1038/msb4100036.

69. Schneider DS. Tracing Personalized Health Curves during Infections. PLOS Biology. 2011;9(9):e1001158. doi:10.1371/journal.pbio.1001158.

70. Kasumovic MM. The multidimensional consequences of the juvenile environment: towards an integrative view of the adult phenotype. Animal Behaviour. 2013;85(5):1049–1059. doi:10.1016/j.anbehav.2013.02.009.

71. Dore AA, McDowall L, Rouse J, Bretman A, Gage MJG, Chapman T. The role of complex cues in social and reproductive plasticity. Behavioral Ecology and Sociobiology. 2018;72(8):124. doi:10.1007/s00265-018-2539-x.

72. Westneat DF, Potts LJ, Sasser KL, Shaffer JD. Causes and Consequences of Phenotypic Plasticity in Complex Environments. Trends in Ecology & Evolution. 2019;34(6):555–568. doi:10.1016/j.tree.2019.02.010.

73. Moller AP, Pomiankowski A. Why have birds got multiple sexual ornaments? Behavioral Ecology and Sociobiology. 1993;32(3):167–176. doi:10.1007/BF00173774.

74. Getty T. The maintenance of phenotypic plasticity as a signal detection problem. The American Naturalist. 1996;148(2):378–385.

75. DeWitt TJ. Costs and limits of phenotypic plasticity: Tests with predator-induced morphology and life history in a freshwater snail. Journal of Evolutionary Biology. 1998;11(4). doi:10.1007/s000360050100.

76. Anderson RM, May RM. Coevolution of hosts and parasites. Parasitology. 1982;85(2):411–426. doi:10.1017/S0031182000055360.

77. Alizon S, Hurford A, Mideo N, Van Baalen M. Virulence evolution and the trade-off hypothesis: history, current state of affairs and the future. Journal of Evolutionary Biology. 2009;22(2):245–259. doi:10.1111/j.1420-9101.2008.01658.x.

78. Mackinnon MJ, Read AF. Genetic Relationships between Parasite Virulence and Transmission in the Rodent Malaria Plasmodium chabaudi. Evolution. 1999;53(3):689–703. doi:10.1111/j.1558-5646.1999.tb05364.x.

79. Mackinnon MJ, Gaffney DJ, Read AF. Virulence in rodent malaria: host genotype by parasite genotype interactions. Infection, Genetics and Evolution. 2002;1(4):287–296. doi:10.1016/S1567-1348(02)00039-4.

80. Mackinnon MJ, Read AF. The effects of host immunity on virulence–transmissibility relationships in the rodent malaria parasite Plasmodium chabaudi. Parasitology. 2003;126(2):103–112. doi:10.1017/S003118200200272X.

81. Paul R, Lafond T, Müller-Graf C, Nithiuthai S, Brey P, Koella J. Experimental evaluation of the relationship between lethal or non-lethal virulence and transmission success in malaria parasite infections. BMC Evolutionary Biology. 2004;4(1):30. doi:10.1186/1471-2148-4-30.

82. Regoes RR, Nowak MA, Bonhoeffer S. Evolution of Virulence in a Heterogeneous Host Population. Evolution. 2000;54(1):64–71. doi:10.1111/j.0014-3820.2000.tb00008.x.

83. Williams PD. New Insights into Virulence Evolution in Multigroup Hosts. The American Naturalist. 2012;179(2):228–239. doi:10.1086/663690.

84. White PS, Choi A, Pandey R, Menezes A, Penley M, Gibson AK, et al. Host heterogeneity mitigates virulence evolution. Biology Letters. 2020;16(1):20190744. doi:10.1098/rsbl.2019.0744.

85. Hawking F, Wilson ME, Gammage K. Evidence for cyclic development and short-lived maturity in the gametocytes of Plasmodium falciparum. Transactions of The Royal Society of Tropical Medicine and Hygiene. 1971;65(5):549–559. doi:10.1016/0035-9203(71)90036-8.

86. Padilla DK, Adolph SC. Plastic inducible morphologies are not always adaptive: The importance of time delays in a stochastic environment. Evolutionary Ecology. 1996;10(1):105–117. doi:10.1007/BF01239351.

87. Hoi AG, Greischar MA, Mideo N. Limited impact of within-vector ecology on the evolution of malaria parasite transmission investment. Frontiers in Malaria. 2024;2. doi:10.3389/fmala.2024.1392060.

88. Mukherjee S, Bassler BL. Bacterial quorum sensing in complex and dynamically changing environments. Nature reviews Microbiology. 2019;17(6):371–382. doi:10.1038/s41579-019-0186-5.

89. Greischar MA, Read AF, Bjørnstad ON. Synchrony in malaria infections: How intensifying within-host competition can be adaptive. American Naturalist. 2014;183(2). doi:10.1086/674357.

90. Chang KH, Tam M, Stevenson MM. Modulation of the Course and Outcome of Blood-Stage Malaria by Erythropoietin-Induced Reticulocytosis. The Journal of Infectious Diseases. 2004;189(4):735–743. doi:10.1086/381458.

91. Mideo N, Barclay VC, Chan BHK, Savill NJ, Read AF, Day T. Understanding and predicting strain-specific patterns of pathogenesis in the rodent malaria Plasmodium chabaudi. American Naturalist. 2008;172(5):214–238. doi:10.1086/591684.

92. Landau I, Boulard Y. Life Cycles and Morphology. In: Killick-Kendrick R, Peters W, editors. Rodent Malaria. Academic Press; 1978. p. 53–84.

93. Gautret P, Miltgen F, Gantier JC, Chabaud AG, Landau I. Enhanced gametocyte formation by Plasmodium chabaudi in immature erythrocytes: Pattern of production, sequestration, and infectivity to mosquitoes. Journal of Parasitology. 1996;82(6). doi:10.2307/3284196.

94. Mideo N, Savill NJ, Chadwick W, Schneider P, Read AF, Day T, et al. Causes of variation in malaria infection dynamics: Insights from theory and data. American Naturalist. 2011;178(6). doi:10.1086/662670.

95. Smith E, Evans G, Foadi J. An effective introduction to structural crystallography using 1D Gaussian atoms. European Journal of Physics. 2017;38(6):065501. doi:10.1088/1361-6404/AA8188.

96. Wood SN. mgcv: GAMs and generalized ridge regression for R. R News. 2001;1.

97. Brancucci NMB, Gerdt JP, Wang C, Niz MD, Philip N, Adapa SR, et al. Lysophosphatidylcholine Regulates Sexual Stage Differentiation in the Human Malaria Parasite Plasmodium falciparum. Cell. 2017;171(7):1532–1544.e15. doi:10.1016/J.CELL.2017.10.020.

98. Regev-Rudzki N, Wilson DW, Carvalho TG, Sisquella X, Coleman BM, Rug M, et al. Cell-Cell Communication between Malaria-Infected Red Blood Cells via Exosome-like Vesicles. Cell. 2013;153(5):1120–1133. doi:10.1016/J.CELL.2013.04.029.

99. Boyle MJ, Wilson DW, Richards JS, Riglar DT, Tetteh KKA, Conway DJ, et al. Isolation of viable Plasmodium falciparum merozoites to define erythrocyte invasion events and advance vaccine and drug development. Proceedings of the National Academy of Sciences of the United States of America. 2010;107(32). doi:10.1073/pnas.1009198107.

100. Stephens R, Culleton RL, Lamb TJ. The contribution of Plasmodium chabaudi to our understanding of malaria. Trends in Parasitology. 2012;28(2). doi:10.1016/j.pt.2011.10.006.

101. Soetaert K, Petzoldt T, Setzer RW. Solving Differential Equations in R: Package deSolve. Journal of Statistical Software. 2020;33(9):1–25. doi:10.18637/jss.v033.i09.

102. Brown PN, Byrne GD, Hindmarsh AC. VODE: A Variable-Coefficient ODE Solver. http://dxdoiorg/101137/0910062. 2006;10(5):p1038–1051. doi:10.1137/0910062.

103. Gerber F, Furrer R. optimParallel: An R Package Providing a Parallel Version of the L-BFGS-B Optimization Method. The R Journal. 2019;11(1):352–358. doi:10.32614/RJ-2019-030.

104. Mullen KM, Ardia D, Gil DL, Windover D, Cline J. DEoptim: An R package for global optimization by differential evolution. Journal of Statistical Software. 2011;40(6). doi:10.18637/jss.v040.i06.

105. Lampinen J, Storn R. Differential Evolution. In: Onwubolu GC, Babu BV, editors. New Optimization Techniques in Engineering. Studies in Fuzziness and Soft Computing. Berlin, Heidelberg: Springer; 2004. p. 123–166. Available from: 10.1007/978-3-540-39930-8_6.

106. Wickham H. ggplot2: Elegant Graphics for Data Analysis. Springer-Verlag New York; 2016. Available from: https://ggplot2.tidyverse.org.

107. R Core Team. R: A language and environment for statistical computing; 2021. Available from: https://www.r-project.org/.

108. Bivand R, Rowlingson B, Diggle P, Petris G, Eglen S. splancs: Spatial and Space-Time Point Pattern Analysis; 2024. Available from: https://CRAN.R-project.org/package=splancs.

